# The Presence of Myelinated Nerves and Schwann Cells in White Adipose Tissue: Proximity to Synaptic Vesicle Containing Nerve Terminals and Potential Role in BTBR *ob*/*ob* Demyelinating Diabetic Neuropathy

**DOI:** 10.1101/2022.08.25.505298

**Authors:** Jake W. Willows, Gilian Gunsch, Emma Paradie, Magdalena Blaszkiewicz, Jeffrey R. Tonniges, Maria F. Pino, Steven R. Smith, Lauren M. Sparks, Kristy L. Townsend

## Abstract

Peripheral neuropathy is a pathophysiological state of nerve degeneration and loss of tissue innervation. The most prominent cause of small fiber neuropathy is diabetes which can be demyelinating in nature, but this has not yet been explored in adipose tissue. Both demyelinating neuropathies and axonopathies implicate Schwann cells (SCs), the peripheral glial required for nerve myelination and regeneration after injury. Here, we perform a comprehensive assessment of SCs and myelination patterns of subcutaneous white adipose tissue (scWAT) nerves, including changes that occur with obesity and other imbalanced energy states in mice and humans. We found that mouse scWAT is densely innervated by both myelinated and unmyelinated sensory and sympathetic nerves. Accordingly, scWAT is home to both myelinating and non-myelinating SCs – the greater proportion of which are myelinating. Furthermore, SCs were found closely associated with synaptic vesicle-containing nerve terminals in scWAT. Obese BTBR *ob/ob* mice exhibit diabetic peripheral neuropathy in scWAT, and display concordant demyelination specific to small fibers, which was also associated with a decrease in the pan-SC marker Sox10 and compensatory increase in Krox20 gene expression. Together this suggests that adipose SCs may be involved in regulating the plasticity or the neuropathy of adipose tissue nerves.

## INTRODUCTION

Peripheral nerves in adipose tissue are demonstrably important for the regulation of tissue function and whole-body metabolic health, as reviewed in [1]. The bidirectional neural communication between adipose and brain is achieved by afferent sensory nerves that communicate from adipose to brain via the dorsal root ganglia and release neuropeptides to adipose tissue when activated, as well as efferent sympathetic nerves that communicate from brain to adipose and release norepinephrine to the tissue. Only recently have the diversity of nerve fibers in adipose tissue, their patterns of innervation, and investigations into nerve terminals or junctions in the parenchyma versus the vasculature been undertaken [2; 3]. From the current work, it is clear that many of the nerves in adipose – whether they are individual sensory or sympathetic axons, or large mixed nerve bundles – are largely myelinated. Additionally, previous scRNAseq studies in adipose have identified populations of Schwann cells (SCs) in the stromal vascular fraction (SVF) [4-6] and SCs have been visualized in visceral white adipose tissue (WAT) [7] and brown adipose tissue (BAT) [3] by TEM. Most recently, neural crest derived SCs were isolated from nerve bundles in mouse and human subcutaneous WAT (scWAT) for use in neurogenic therapies [8]. Taken together, more knowledge is needed to better understand the contribution of adipose tissue SCs to nerve function and tissue function.

Peripheral neuropathy, or the dying-back of peripheral axons innervating tissues and organs, can cause severe pain and discomfort and blunts tissue neural communication with the brain, leading to numerous health complications [9; 10]. We previously demonstrated that aging leads to peripheral neuropathy in scWAT [11], and that in obese and diabetic BTBR mice homozygous for the spontaneous *Lep*^*ob*^ mutation (BTBR *ob/ob*) [12] the peripheral neuropathy also extended to metabolically-relevant tissues like adipose and muscle [11]. Deletion of brain derived neurotrophic factor (BDNF), a neurotrophic factor implicated in human obesity, from myeloid-lineage neuroimmune cells also causes a ‘genetic denervation’ of adipose tissue [13] that is accompanied by worsened metabolic phenotype. Small fiber peripheral neuropathy is common with altered metabolic disease states such as diabetes or aging, and these may involve demyelinating neuropathies or reduced axon outgrowth due to impaired SC function in nerve repair and regeneration [14]. From our prior work, small fiber innervation in scWAT may be particularly important for metabolic control [13].

SCs support normal nerve physiology through the formation of myelin sheaths, which enable electrical conductance by providing insulation for the axon’s ionic flow. Each myelinating SC (mSC) surrounds a single axon and several mSCs are required to myelinate the entire length of an axon. Non-myelinating SCs (nmSCs) are usually associated with smaller diameter axons (Remak bundles), such as many sensory axons and axons of the autonomic nervous system, as reviewed in [14]. Therefore, demyelination of nerves results in reduced nerve conductance and loss of functions (motor and sensory deficits, for example). The unique capability for plasticity and regeneration of nerves in the peripheral nervous system (PNS) is due largely to the ability of SCs to respond to and repair damaged nerves by phenotypically switching to “repair SCs” [15].

The conversion, or transdifferentiation, of mature SCs to repair SCs involves down-regulation of myelin-production genes (such as myelin basic protein (MBP), Krox20, and myelin protein zero (MPZ)), and upregulation of glial fibrillary acidic protein (GFAP), the neurotrophic factor receptor p75NTR, and c-Jun [16]. Restoration of c-Jun expression in SCs is able to rescue failed nerve regeneration in aging mice [17], and is thus considered a master regulator of repair SC phenotypic conversion. The exact signals from damaged nerves (released during a process known as Wallerian degeneration) that promote SC conversion to repair SCs is currently unknown. Given the striking plasticity of SCs in response to their local environment, it is likely that SC phenotypes and activity differ by tissue and by tissue state [14]. Therefore, a better characterization of adipose-resident SCs is warranted to further our current understanding of SC plasticity and their role in adipose-related pathologies, including diabetic neuropathy

## RESEARCH DESIGN AND METHODS

### Experimental model and subject details - Mice

All mice were obtained from The Jackson Laboratory (Bar Harbor, ME) and/or bred at our mouse facilities at University of Maine and The Ohio State University. Animals were housed in a temperature-controlled environment, kept on a 12 hr light-dark cycle, and allowed access to food and water *ad libitum* (unless stated otherwise for a particular study). For all studies animals were euthanized using CO_2_ followed by cervical dislocation. All procedures were performed in compliance with the National Institute of Health Guide for the Care and Use of Laboratory Animals and were approved by an Institutional Animal Care and Use Committee.

#### PGP9.5-EGFP^+/-^ Reporter Mice

Male and female PGP9.5-EGFP^+/-^ (*C57BL/6-Tg(Uchl1-EGFP)G1Phoz/J*, Stock # 022476) pan-neuronal reporter mice were used for microscopy experiments to investigate adipose innervation. Mice were housed 2-5 a cage, fed a standard chow diet and were aged 10-26 weeks prior to tissue collection.

#### Sedentary/Exercised

Adult (12-15 week old) C57BL/6J male mice, age and body weight matched, were assigned to either sedentary or exercised groups. Animals were single housed in running wheel cages that allowed *ad libitum* access to running for a period of 7 days. Wheel running was monitored using odometers (magnets were placed on the outer arm of the running wheels with the sensor attached to the inside of the cage). Sedentary (control) animals were single caged with locked running wheels for the same period.

#### BTBR *ob/ob* mice

BTBR *+/+* (WT) and BTBR *ob/ob* (MUT) mice (BTBR.Cg-*Lep*^*ob*^*/*WiscJ) were fed a standard chow diet, housed two to a cage, and aged at least 12 weeks, when they exhibit a strong phenotype (including obesity and diabetes) for all experiments.

#### Cold exposure experiments

Adult (8 week old) male C57BL/6J mice were housed two to a cage, and either maintained at room temperature (RT), continuously cold exposed (at 5ºC), or kept at thermoneutral temperature (30ºC) for 3 days with *ad libitum* access to food and water. All cold exposure experiments occurred in a diurnal incubator with 12 hr light/dark cycle and humidity control (Caron, Marietta, OH, USA).

#### Diet-induced obesity

Adult (11 week old) male C57BL/6J mice, housed 2-3 to a cage, were fed either chow control or a 58% high fat diet (HFD) (Research Diets Cat #D12330) *ad libitum* for 19 weeks. Animals were euthanized with CO_2_ and inguinal scWAT was collected and frozen before being processed for qPCR.

#### Aging

C57BL/6J male mice were housed 2-4 in a cage and aged to 15 weeks (young) and 75 weeks (aged). Tissues where snap-frozen in liquid nitrogen upon collection for gene expression analysis using RT-qPCR. For ageing related FACS studies, C57BL/6J male mice were aged to 4 months, 8 months, and 15 months.

### Experimental model and subject details – Human tissues

Human abdominal scWAT were obtained from 23 individuals, 11 were healthy lean (BMI ≤ 25) and 12 were healthy obese (BMI >30) individuals who participated in a cross-sectional study at the Translational Research Institute at AdventHealth. The study protocol was approved by the AdventHealth Institutional Review Board and was in line with the Declaration of Helsinki. Samples were collected from informed and consenting male (lean N=2, 18-23 years old; obese N=3, 35-64 years old) and female (lean N=9, 23-49 years old; obese N=9, 25-55 years old) subjects during elective surgery. Abdominal scWAT was obtained under local anesthesia [18]. Tissue was immediately snap-frozen in liquid nitrogen and then processed for RNA extraction. RNA was used to measure gene expression by real-time qPCR

### Whole mount immunofluorescence (IF)

Whole inguinal and axillary scWAT depots were carefully removed to remain fully intact and fixed in 2% PFA (Sigma, Cat#P6148) for 16hrs (or 24hrs for obese BTBR *ob/ob*) at 4°C. Tissues were Z-depth reduced and subsequently blocked in 2.5% BSA / 1% Triton X-100 / 1X PBS for 24hrs at 4°C and incubated in 0.1% Typogen Black (Sigma, Cat#199664) to reduce autofluorescence. BTBR *ob/ob* tissues did not receive autofluorescence quenching. Tissues were incubated in primary antibody solution for 48hrs at 4°C, rinsed in 1X PBS, and incubated in secondary antibody solution for 24hrs at 4°C. These steps were repeated for additional co-labeling. When desired, tissues were incubated in 100 ng/mL DAPI (Sigma-Aldrich, Cat#D9564) for 1hr at room temperature to label nuclei. Finally, tissues were mounted on glass slides and imaged. For additional details see [2] and the accompanying peer-reviewed protocol [19].

### Confocal microscopy

Confocal micrographs were captured on a Leica Stellaris 5, laser scanning confocal microscope using LASX software. Fluorescent labels were excited with either a diode 405 nm laser (DAPI, Autofluorescence) or a white light laser: EGFP (499 nm), Alexa Fluor 555 (553 nm), Alexa Fluor Plus 594 (590 nm), Alexa Fluor Plus 647 (653 nm). Emission spectra were tuned specifically for each fluorophore or groups of fluorophores to reduce and eliminate crosstalk. Multiple channels were scanned sequentially, and all channels were line averaged 3 times. Photons were detected with Power HyD S detectors. Objectives included: HC PL APO 10x/0.40 CS2, HC PL APO 40x/1.30 OIL CS2, and HC PL APO 63x/1.40 OIL CS2. PinholeAiry 1.00 AU. Confocal zoom was applied to further increase magnification when necessary. Entire tissues were visually scanned, and representative images were captured at 2048×2048 pixel resolution as Z-stacks (1-6 µm step size) that were maximum intensity projected. LUTs were adjusted to improve structure visualization. Digital cross-sections were captured by utilizing the XZY scan function. One iteration of a 5-kernel median noise filter was applied. Image processing performed in Leica LASX software.

#### Deconvolution

In specific instances (stated in figure legends) images were acquired using Lightning (Leica) 3D deconvolution. Z-stack images were captured at Nyquist lateral and axial resolutions at 63X objective magnification with a PinholeAiry of 0.50 AU with varying confocal zooms applied. Adaptive deconvolution was calculated for a 1.4429 refractive index. Deconvolved z-stacks were maximum intensity projected and LUTs were adjusted to improve structure visualization. Image processing performed in Leica LASX software.

#### Epifluorescence microscopy

Epifluorescence micrographs were captured on a Nikon Eclipse E400 epifluorescence microscope using a Hamamatsu ORCA-Flash4.0 V2 Digital CMOS monochrome camera. Alexa Fluor 555 fluorophores were excited using a Cy3 filter cube. Objectives used: Nikon CFI Plan Fluor 20x/0.50 and Nikon CFI Plan Fluor 40x/0.75. Images were captured utilizing the extended depth of field (EDF) function; LUTs were adjusted to improve structural visualization. Post processing was performed in Nikon Elements BR software.

### Neuromuscular junction immunostaining

Medial gastrocnemius muscle was fixed in 2% PFA for 2hrs and teased apart. Tissue was blocked in 2.5% BSA / 1% Triton X-100 / 1X PBS for 24hrs at 4°C. Tissues were incubated in a series of primary and secondary antibody solutions each lasting 24hrs and performed at 4°C on a rotator. Tissues were washed in 1X PBS between incubations. Primary antibodies SV2 and 2H3 were incubated simultaneously with α-bungarotoxin (BTX) conjugated to Alexa Fluor 555 (1 mg/mL, Thermofisher, Cat#B35451) to label pre- and post-synapse. This was followed by the secondary antibody goat anti-mouse IgG1 Alexa Fluor 488, and then the primary antibody MPZ followed by its secondary antibody goat anti-rabbit IgG Alexa Fluor 647 Plus. Finally, the tissues were incubated in 100 ng/mL for 1hr at room temperature to label nuclei and mounted on glass slides for imaging.

### Antibodies used for immunostaining

#### Primary antibodies

MPZ (1:250, Abcam, Cat#ab31851); SOX10 (1:200, Abcam, Cat#ab227680); CGRP (1:200, EMD Millipore, Cat#PC205L); TH (1:250, EMD Millipore, Cat#AB152); NCAM (1:250, Millipore, Cat#AB5032); SV2 (1:250, Developmental Studies Hybridoma Bank, Cat#SV2); 2H3 (1:500, Developmental Studies Hybridoma Bank, Cat#2H3); SYN1 (1:200, Cell Signalling, Cat#5297); TUBB3 conjugated to Alexa Fluor 488 (1:200, Abcam, Cat#ab195879); MBP conjugated to Alexa Fluor 555 (1:500, Cell Signaling, Cat#84987).

#### Secondary antibodies

Goat anti-Rabbit IgG (H+L) Cross-Adsorbed Secondary Antibody, Alexa Fluor 555 (1:500, Thermofisher, Cat#A-21428); Goat anti-Rabbit IgG (H+L) Highly Cross-Adsorbed Secondary Antibody, Alexa Fluor Plus 594 (1:1000, Thermofisher, Cat#A32740); Goat anti-Rabbit IgG (H+L) Highly Cross-Adsorbed Secondary Antibody, Alexa Fluor Plus 647 (1:250, Thermofisher, Cat#A-32733), Goat anti-Mouse IgG1 Cross-Adsorbed Secondary Antibody, Alexa Fluor 488 (1:500, Thermofisher, Cat#**A-21121)**.

### Adipose stromal vascular fraction (SVF) dissociation

SVF from bilateral whole inguinal adipose depots was isolated as previously described [12]. Inguinal scWAT was quickly harvested and minced in DMEM (high glucose, serum free, pre-warmed in a 37ºC water bath) containing 2 mg/mL of Collagenase A (Sigma-Aldrich, Cat#10103586001) at a volume of 20 mL/g of tissue. Collagenase was added to warmed media immediately before tissue dissection. Inguinal scWAT was pooled bilaterally from each animal for FACS analysis of exercise and ageing cohorts. For FACS analysis at basal state, bilateral depots from 4 animals were pooled together after the dissociation step. Minced tissue in dissociation media was placed in 50 mL conical tube and transferred to a shaking warm water bath (90 rotations/min at 37ºC). Dispersion of cells was furthered via gentle vortexing and trituration using Pasteur Pipettes at various bores until full dissociation was achieved and floating adipocytes were visible. Samples were filtered through 100 µM cell strainers and rinsed with DMEM then centrifuged at 500 g for 10 min to separate adipocytes and SVF pellet. SVF pellet was incubated with 500 µl of red blood cell lysis buffer for 2 min on ice. Lysis was stopped by the addition of 2 mL of DMEM containing 5% FBS. Cells were centrifuged at 500 g for 5 min at 4°C and resuspended in 100 µL of FACS buffer (1X PBS with FBS and EDTA) for cell sorting.

### Fluorescence-activated cell sorting (FACS)

For antibody labeling, 2 µL FACS block (BSA and FBS in 1X PBS) was added to SVF resuspended in FACS buffer (as described above) and then left to sit for 15-20 min on ice, after which 500 µL FACS buffer was added, and samples were spun at 1800 rpm for 5 min. Samples were resuspended in 100 µL FACS buffer with conjugated antibodies. Antibodies used included; Anti-CD45-BV421 (Biolegend, Cat. # 103133 (clone 30-F11)), Anti-O4-APC (Miltenyi Biotec, Clone O4), Anti-p75-VioBright FITC (Miltenyi Biotec, Clone REA648). Cells were washed 1-2 times by centrifugation at 1500 rpm for 5 min and then resuspended in 300 µL of FACS buffer. UltraComp eBeads (Invitrogen # 01-2222-42) were used for compensation controls. DAPI exclusion was used for viability. Sorting was performed on a BD™ FACS Aria II™ cell sorter or BD™ Influx equipped with a 100uM nozzle to accommodate Schwann cell size. Cells were gated on FSC/SSC, live cells and CD45 to exclude immune cells. Cells representing Schwann cell populations (CD45-p75+, CD45-p75+O4+, or CD45-O4-), were sorted into 400 µL of Trizol (Zymo, Irvine, CA, USA; Cat. #R2050-1-200). FACS performed sorts reported in this paper were performed at The Jackson Laboratory in Bar Harbor, ME and at The Nationwide Children’s Hospital Flow Cytometry Core Facility in Columbus, OH.

### RNA extraction and real-time quantitative PCR (qPCR)

Zymo DirectZol RNA extraction kit (Zymo, Irvine, CA, USA; Cat. #R2052) was used for RNA extraction from whole tissue and the RNeasy Micro Kit (Qiagen, Cat. # 74004) was used to isolate RNA from sorted cells. RNA yield was determined using a Nanodrop and cDNA was synthesized using High-Capacity Synthesis Kit (Applied Biosystems, Foster City, CA, USA; Cat. #4368813). Real-time quantitative polymerase chain reaction was performed using SYBR Green (Bio-Rad, Cat#1725271) on a CFX384 real-time PCR detection system (Bio-Rad, Hercules, CA, USA). Gene expression was normalized to housekeeper gene *Ppia* and *Sox10* for analysis. Primers used for qPCR are listed in Supplemental Table 1.

### Western blot (WB)

Protein expression was measured by western blotting analysis of scWAT lysates. Whole adipose depots were homogenized in RIPA buffer with protease inhibitors in a Bullet Blender. A Bradford assay was performed to measure total protein from which equal concentrations of protein lysates were prepared in Laemmli buffer using 1X PBS as diluent. 30 µg of protein were loaded per lane of a 10% polyacrylamide gel, and following gel running, proteins were transferred to PVDF membrane and incubated with 10% Roche Blocking Reagent for 1 hr at room temperature prior to antibody incubation. The membrane was bisected horizontally at 37 kDa to avoid the need for stripping proteins later. Primary antibodies included: TH (62 kDa, 1:250, EMD Millipore, Cat#AB152); SOX10 (49 kDa, 1:400, Abcam, Cat#ab155279); MPZ (25 kDa, 1:1000, Abcam, Cat#ab31851); β-Tubulin (55 kDa, 1:1000, Cell Signaling Technology, Cat#2146BC); Cyclophilin B (21 kDa, 1:40,000, Abcam, Cat#16045). Primary antibodies were incubated overnight at 4°C on a rotator with gentle agitation. Membranes were rinsed with 1X TBS-T and then incubated in anti-rabbit HRP-linked secondary antibody (1:3000, Cell Signaling Technology, Cat#7074) for 1hr a room temperature. Blots were visualized with enhanced chemiluminescence (ECL; Pierce) on a Syngene G:BOX. TH and SOX10 expression were normalized to Cyclophilin B, and MPZ was normalized to β-Tubulin and quantified by densitometry in Fiji [19].

### Statistical analyses

For all animal experiments, mice were body weight matched and then randomized to experimental or control groups to mitigate differences in starting body weights. ROUT outlier test (Q=1%) was performed on raw data to identify and remove statistical outliers within data sets. FACS data was analyzed by two-tailed Student’s t-test or two-way ANOVA with Tukey’s multiple comparison test for mixed models. qPCR data was analyzed for each gene individually by either unpaired two-tailed Student’s t-test or one-way ANOVA with multiple comparisons. Linear regression analysis was performed with Goodness of Fit measured by R-squared and the significance of slope determined by F-test, when comparing two linear regressions statistical difference between slopes and Y-intercepts were analyzed. Western blot was analyzed by unpaired Student’s t-test. G-ratio was analyzed by unpaired Student’s t-test. Neurite density was analyzed as unpaired Student’s t-test. All error bars are SEMs. Statistical calculations for determining significance were calculated in GraphPad Prism software (La Jolla, CA, USA). For all figures statistically significant p-values are displayed on each graph. n. s. = not significant.

### BTBR ob/ob scWAT nerve bundle processing, imaging, and quantification

A prominent nerve bundle traversing into the inguinal scWAT depot was excised from each mouse and samples were fixed in a 2% PFA / 2.5% glutaraldehyde mixture, postfixed with 1% osmium tetroxide, and then en bloc stained with 1% aqueous uranyl acetate. The samples were dehydrated in a graded series of ethanol and embedded in Eponate 12 epoxy resin (Ted Pella Inc., Redding, CA). One-micron thick sections were cut with a Leica EM UC7 ultramicrotome (Leica microsystems Inc., Deerfield, IL) and stained with toluidine blue. Images of nerve bundle cross-secitons were acquired with a Zeiss Axioskop microscope (Carl Zeiss Microscopy, LLC, White Plains, NY) using Zeiss Achroplan 20X/0.45 Ph2 and Zeiss Plan-Apochromat 63X/1.4 Oil Ph3 objective lenses. Images were captured with a Nuance multispectral imaging camera (PerkinElmer, Waltham, MA).

Intact whole axillary scWAT depots were excised and immunostained for TH. Alexa fluor plus 594 was excited at 590 nm, and emitted photons were detected for 600-700 nm. Laser intensity (10%) and detector gain (8%) remained constant for all images. The entirety of each whole tissue was visually scanned at 10X objective magnification (1.00 confocal zoom) and a representative 5×5 tiled (22.85 mm^2^) area was captured as a series of Z-stacks (10 µm step size) extending through the full thickness of each tissue (120-230 µm) and were maximum intensity projected. Images were processed for quantification in Fiji [20] by first applying background subtraction (50 pixel rolling ball radius). Next, a threshold (30-255) was applied to further remove autofluorescent background. Area of remaining pixels was measured.

### Data and resource availability

The datasets generated during and/or analyzed during the current study are available from the corresponding author upon reasonable request. No applicable resources were generated or analyzed during the current study.

## RESULTS

### Patterns of nerve myelination in scWAT

We used C57BL/6-Tg(Uchl1-EGFP)G1Phoz/J mice which endogenously express a pan-neuronal GFP reporter (henceforth referred to as PGP9.5-EGFP^+/-^) to investigate the relative proportions of myelinated and non-myelinated nerves in inguinal scWAT (Figure 1A-C). To label the myelin sheath of adipose-resident peripheral nerves, we used antibodies against two myelin specific proteins, MPZ that is more commonly found in the PNS, and MBP that is more commonly found in the CNS but also localized to PNS myelin [21]. Co-expression of PGP9.5-EGFP^+/-^ with either MPZ or MBP was interpreted as a myelinated fiber and the absence of either marker on a PGP9.5-EGFP^+/-^ nerve was interpreted as a non-myelinated fiber. Each whole adipose tissue depot was systematically visually scanned for qualitative analyses, and representative images for each tissue were captured of nerve bundles, nerves innervating blood vessels, and nerves in the parenchyma.

**Figure 1:**
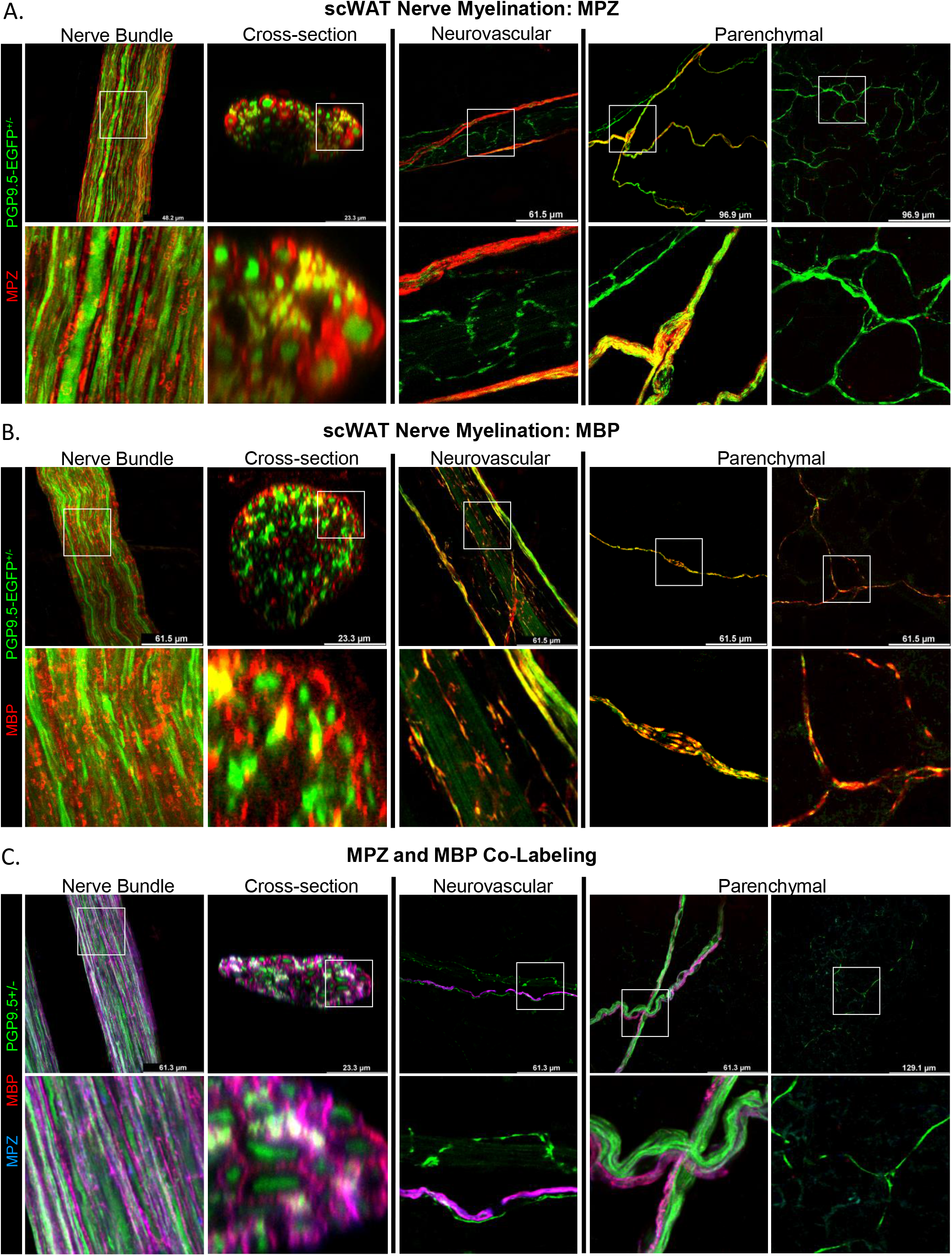
Immunofluorescence in subcutaneous white adipose tissue (scWAT) reveals a heterogenous mix of myelinated and non-myelinated axons. Intact inguinal scWAT depots were excised from PGP9.5-EGFP^+/-^ reporter mice and immunolabeled for myelin with MPZ and MBP. Representative images are displayed of nerve bundles (top-down and cross-sectional), nerves interacting with blood vessels, and those in the parenchyma. Each marker was immunostained and analyzed individually **(A-B)** and co-labeled **(C)** to investigate co-expression. Structures co-labeled with PGP9.5-EGFP^+/-^ (green) and either of the myelinating markers (red) were identified as myelinated nerves **(A-B)**. Overlap of MPZ (red) and MBP (blue) was used to determine extent of MPZ and MBP co-expression **(C)**. White boxes show select regions digitally enlarged to aid visualization. Images were captured on Stellaris 5 confocal microscope. See Supplemental Figure S1 for single-color channels of each image.

All large nerve bundles (>25 μm diameter when viewed from above) in scWAT presented with myelination, as indicated by MPZ/MBP labeling (Figure 1A and 1B, Nerve Bundle). These large bundles displayed numerous branching points that eventually led to individual myelinated fibers that branched further into thinly or unmyelinated fibers. The presence of myelin on each nerve bundle appeared unevenly distributed, and myelin sheaths could not be traced along each individual axon in the field of view. Digital cross-sectioning of these nerve bundles provided a clear view of individual axons within the bundle, and the MPZ+ and MBP+ myelin sheaths around them (Figure 1A and 1B, cross-section). In this view it was evident that MPZ and MBP labeling were specific to the myelin sheath. Light scatter and a potential inability for antibodies to penetrate the full diameter of nerve bundles resulted in the superficial aspect (closest to the light source) to appear brighter than the nerve fibers distal from the light source, and was partially obscured by the overlying nerves within the bundle. Myelin was present throughout the depth of each bundle, but localization of MPZ+ and MBP+ staining to individual fibers was most obvious superficially.

Nerves innervating the adipose vasculature followed consistent patterning to what has been described previously [2]; small unmyelinated fibers contacting vessel walls with myelinated fibers traversing in parallel with the vessel. Here we note surprising variation between the markers for myelin sheaths (Figure 1A and 1B, neurovascular). MPZ consistently labeled one or two myelinated fibers running parallel to most blood vessels, excluding capillaries, and the small fibers connecting with the vessel wall were MPZ-, supporting previous findings [2] that the nerves directly innervating tissue vasculature are unmyelinated sympathetic nerves (Figure 1A, neurovascular). By contrast, MBP staining of neurovascular fibers was less consistent. MBP was observed to label the thicker nerves running parallel to vessels as well as many of the smaller fibers contacting the vessel wall (Figure 1B, neurovascular). As with neurovascular labeling, MPZ was expressed only on a subset of nerve fibers in the parenchyma with the majority being MPZ- (Figure 1A, Parenchymal). MBP seemed to label the parenchymal fibers found alongside adipocytes with less discrimination (Figure 1B, Parenchymal).

Because MPZ and MBP had slightly different patterns of labeling, it was necessary to co-stain with these antibodies to gauge the extent of their overlap (Figure 1C). Co-labeling MPZ with MBP demonstrated complete co-expression in the myelin sheaths around nerve fibers within bundles (Figure 1C, nerve bundle, cross-section). Although MBP was observed to label a greater number of small fibers, co-labeling with MPZ showed that in large part, labeling around vasculature (Figure 1C, Neurovascular) and in the parenchyma (Figure 1C, Parenchymal) was consistent for both markers.

### Distinguishing sympathetic and sensory nerve myelination in scWAT

Given scant prior reports of myelination status for the sensory and sympathetic innervation in WAT, we assessed myelin presence on sympathetic nerves that were labeled for tyrosine hydroxylase (TH), the rate-limiting enzyme for norepinephrine synthesis, as these nerves comprise the majority of scWAT innervation [22]. We also labeled sensory nerves for the sensory-specific neuropeptide Calcitonin Gene-Related Peptide (CGRP). This may under-represent the entirety and diversity of adipose sensory nerves since neuropeptide diversity and non-peptidergic sensory fibers are only beginning to be described in scWAT [2].

Intact inguinal scWAT depots were excised from PGP9.5-EGFP^+/-^ mice and co-stained for TH and MBP. TH+ axons comprised only a subset of the axons within each myelinated nerve bundle (Figure 2A, nerve bundle, cross-section). Neurovascular axons were almost entirely TH+, excluding rarely observed myelinated nerves running in parallel with blood vessels, which were TH-/MBP+ (Figure 2A, neurovascular). We observed that the majority of TH+ fibers co-expressed MBP around vasculature and in the parenchyma (Figure 2A, parenchymal).

**Figure 2:**
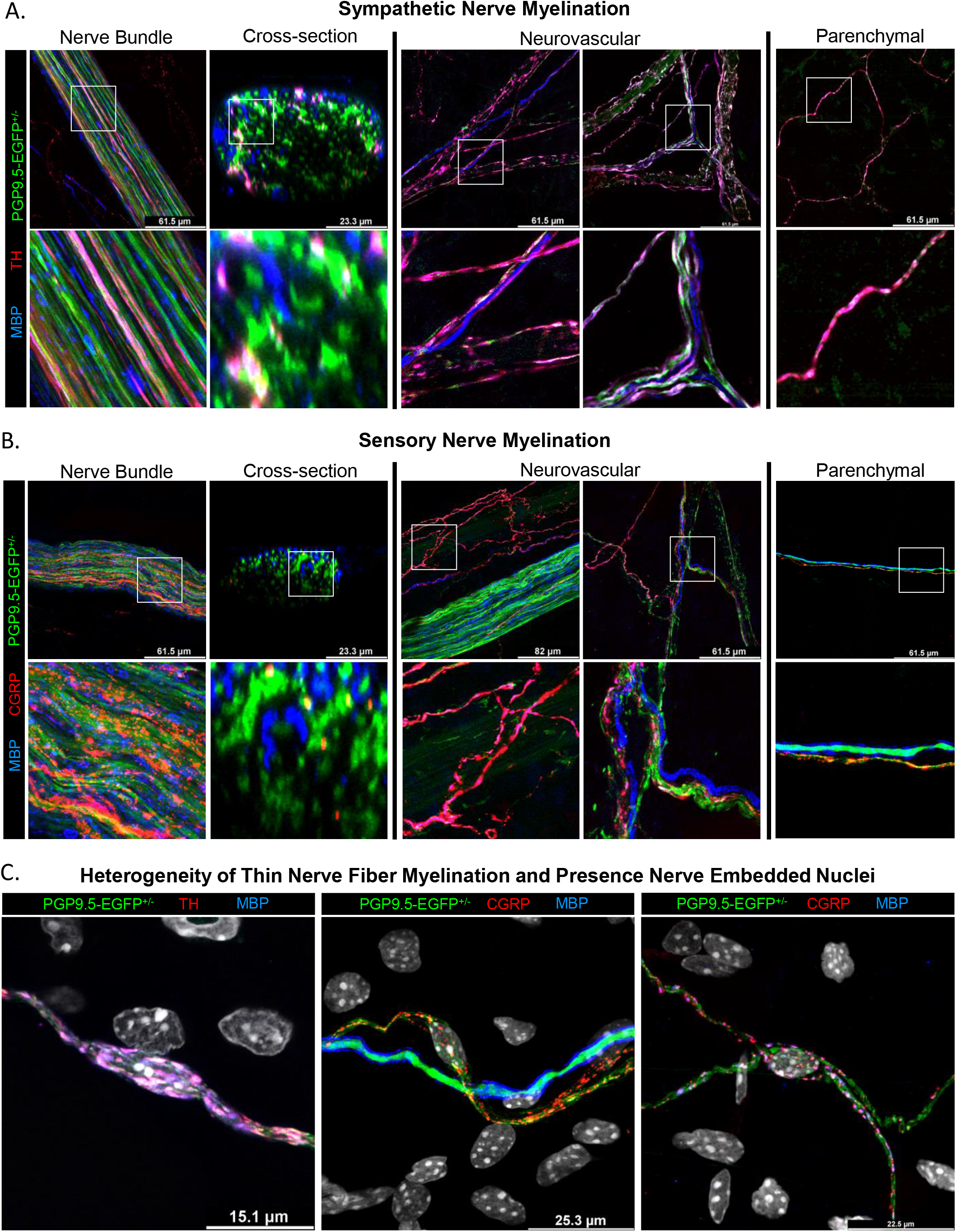
Sympathetic and sensory nerve myelination in scWAT. To investigate the status of sympathetic and sensory nerve myelination, intact inguinal scWAT depots were excised from PGP9.5-EGFP^+/-^ (green) reporter mice and co-labeled for MBP (blue) and either the sympathetic nerve marker TH (red) **(A)** or the sensory nerve marker CGRP (red) **(B)**. Representative images of nerve bundles, vascular innervation, and parenchymal innervation are displayed. White boxes were digitally enlarged to aid in visualization. Red and blue overlap identified myelinated sympathetic **(A)** and Sensory nerves **(B)**. Axons in the parenchyma demonstrate nuclei (DAPI, grey) embedded in both sympathetic and sensory nerves **(C)**. Images were captured on Stellaris 5 confocal microscope. See Supplemental Figure S2 for single-color channels of each image.

PGP9.5-EGFP^+/-^ nerve bundles co-stained for CGRP and MBP revealed punctate CGRP+ labeling across the length of a subset of the axons within myelinated nerve bundles (Figure 2B, nerve bundle). Digital cross sections supported this finding, although due to the punctate nature of the staining (likely because the vesicles containing CGRP were not uniformly distributed along the axon), it is possible that digital cross sections underrepresented the total number of CGRP+ axons in each bundle (Figure 2B, cross-section). Neurovascular imaging of CGRP+ nerves showed that the majority of CGRP+ axons outside of nerve bundles were closely aligned with blood vessels (Figure 2B, neurovascular). This was not surprising given CGRP’s known role in vasodilation [23; 24]. Small axons in contact with the vessel wall were largely CGRP-/MBP-, and only rarely were CGRP+/MBP+ (Figure 2B, neurovascular). Most parenchymal nerves were CGRP-/MBP-, with infrequent CGRP+/MBP-, CGRP+/MBP+, and CGRP-/MBP+ axons also present (Figure 2B, Parenchymal). Parenchymal nerve fibers presented with what appeared to be cell bodies residing between axons and forming gaps between adjacent fibers (Figure 2C), which were presumed to be Schwann cells or neuroimmune cells since neuronal cell bodies are in the dorsal root ganglia and not in the tissue. DAPI labeled nuclei were found embedded in both myelinated (MBP+) and unmyelinated (MBP-) axons, regardless of CGRP or TH expression (Figure 2C). Combined, these data exemplify the heterogeneity of nerve fibers within scWAT including myelinated sensory and sympathetic axons.

### SCs in scWAT

To confirm that the nuclei residing between adjacent nerve fibers and around scWAT axons were SCs, we labeled whole inguinal scWAT from PGP9.5-EGFP^+/-^ reporter mice for the SC lineage-determining transcription factor SOX10 [25; 26], which was used as a pan-SC marker. DAPI+ nuclei residing between adjacent fibers and along the parenchymal nerves were found to be SOX10+ (Figure 3A), thus confirming their SC identity. A thorough analysis of the whole tissue revealed that SOX10+ cells were prominently distributed throughout the tissue (Figure 3B), with the majority localized to nerve bundles. SOX10+ cells contributed to most of the DAPI labeled nuclei within and around each nerve bundle (Figure 3B, nerve bundle, cross-section). The remainder of the nuclei associated with each bundle are speculated to be neuroimmune cells, based on prior data [13]. The second greatest proportion of SOX10+ cells were observed in contact with the nerves innervating tissue vasculature (Figure 3B, neurovascular), and relatively few SOX10+ cells were observed in the parenchyma around adipocytes (Figure 3B, parenchymal). Importantly, the SOX10 antibody used in this manuscript also demonstrated off-target labeling of what appeared to be mammillary ducts and/or lymphatic vessels (Supplemental Figure S3A). Fortunately, SCs and ducts/vessels were easily distinguished from one another due the distinct morphology of cells versus vessels (Supplemental Figure S3B).

**Figure 3:**
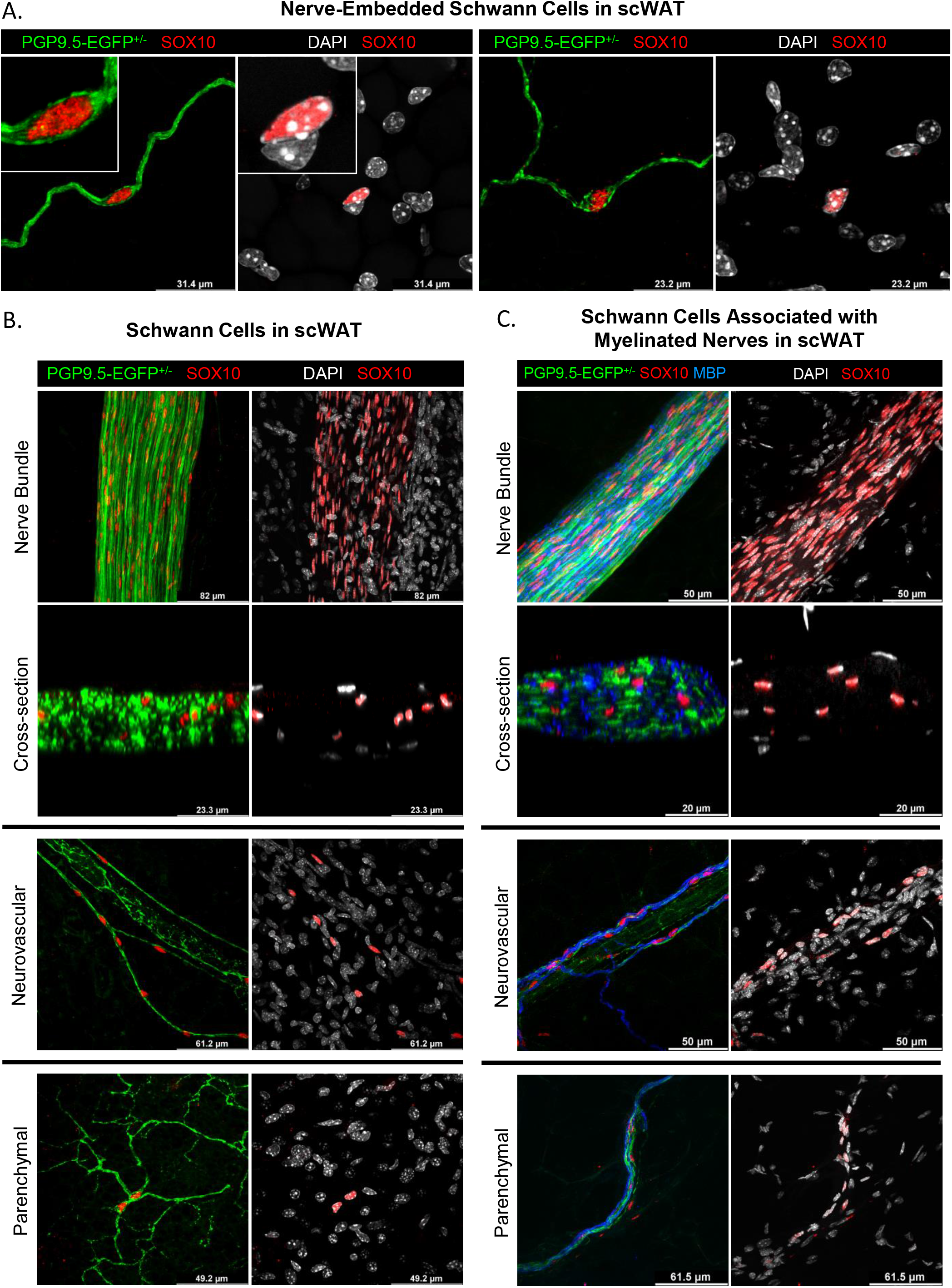
Schwann cell (SC) bodies are embedded in heterogenous nerves within scWAT. Intact inguinal scWAT depots were excised from PGP9.5-EGFP^+/-^ (green) reporter mice and co-stained to label SCs (SOX10, red) and nuclei (DAPI, grey) **(A-B)**. High magnification image of SOX10 positive nuclei embedded in small parenchymal nerve fibers **(A)**. Representative images of SOX10 positive nuclei distribution throughout scWAT **(B)**. SC localization in relation to myelinated nerves stained with MBP **(C)**. Overlap of red and grey identified SCs **(A-C)**. Overlap of green and blue identified myelinated nerves **(C)**. Images were captured on Stellaris 5 confocal microscope. See Supplemental Figure S3 for single-color channels of each image.

Although SOX10 staining cannot differentiate between mSCs and nmSCs, we observed that the greatest proportion of SOX10+ cells were distributed in structures that we had previously observed to have significant myelination [2]. To confirm this, we co-stained scWAT from PGP9.5-EGFP^+/-^ reporter mice for SOX10 and MBP (Figure 3C). As expected, most SOX10+ cells were associated with MBP+ nerves, indicating they were mSCs. There were also SOX10+ cells associated with unmyelinated nerves in the parenchyma, but these were less common (Figure 3C).

#### The neuro-adipose nexus (NAN) is preceded by SCs

Following the recent discovery of putative nerve terminals in scWAT, a structure where axons terminate by wrapping around individual adipocytes which we termed the ‘neuro-adipose nexus’ (NAN) [2], we sought to learn more about these structures by labeling for various nerve, synaptic, and SC markers (Figure 4). Whole inguinal scWAT depots were excised from C57BL/6J mice and immunostained for sympathetic nerves (TH). Tissue autofluorescence was captured intentionally to visualize adipocyte boundaries and tissue structure along with TH labeling (Figure 4A), as is often exploited in tissue clearing experiments. Images were also captured with TH labeling that did not include tissue autofluorescence (Supplemental Figure S4A), as control. NANs were labeled clearly by TH and were visualized in clusters (2-4 adipocytes) as well as individually (Figure 4A, Supplemental Figure S4A). These images revealed that branching axons terminate on the cell surface of adipocytes (Figure 4A). Axons forming each NAN were pearled by varicosities, similar to those observed in autonomic neuroeffector junctions [27] (Figure 4A).

**Figure 4:**
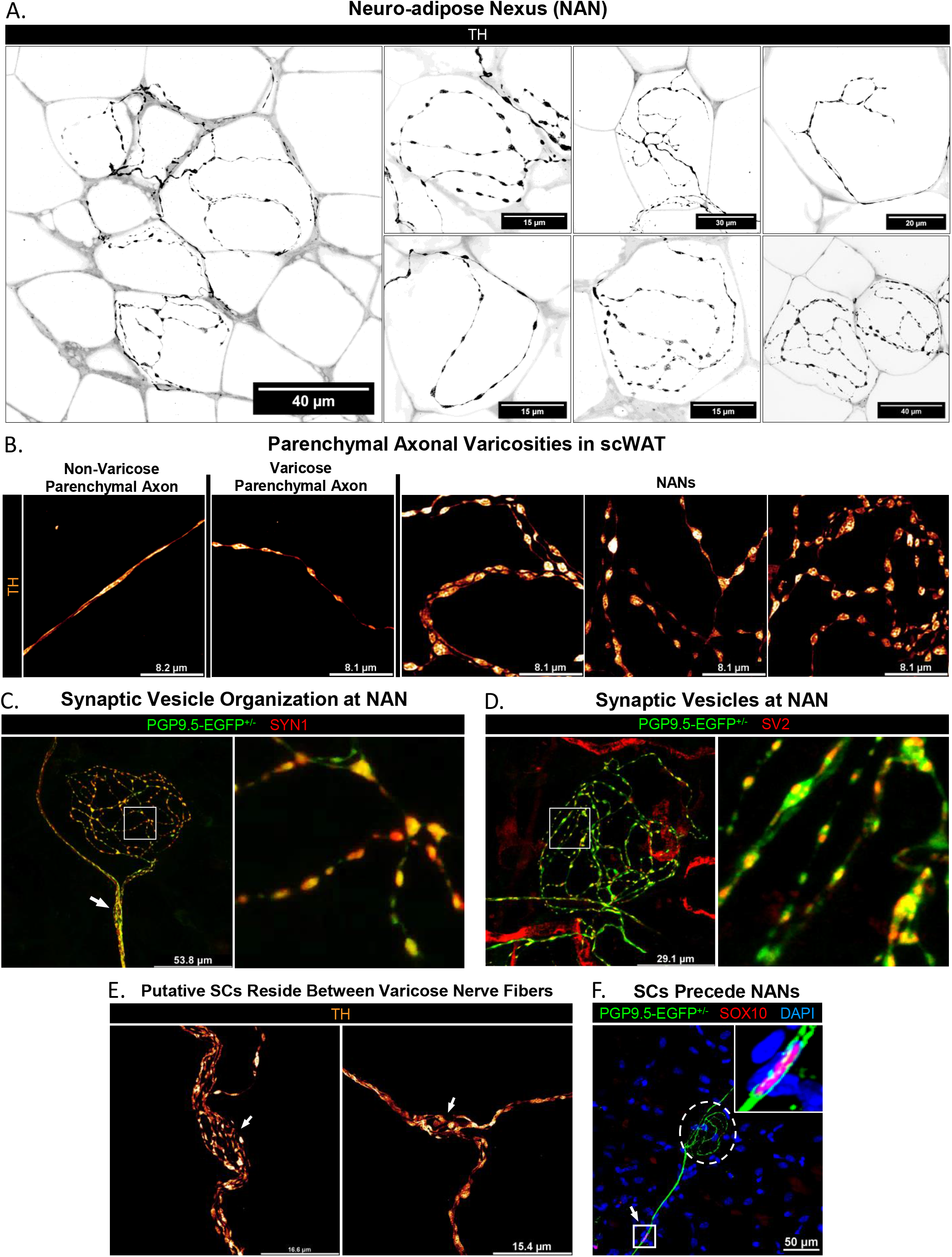
The neuro-adipose nexus (NAN) is densely pearled by synaptic vesicle containing axonal varicosities and is often immediately preceded by SCs. Intact inguinal scWAT depots were excised from C57BL/6J mice and labeled for TH which revealed NANs throughout the tissue parenchyma terminating on single adipocytes or small clusters of adipocytes **(A)**. TH labeling of NANs were merged with single Z-planes of purposefully captured tissue autofluorescence to display adipocyte boundaries in relation to the spread of axons in each NAN; displayed as inverted monochrome images **(A)**. TH+ axons are black overlying white adipocytes with grey extracellular matrix separating each cell **(A)**. NANs were characteristically densely populated by axonal varicosities when compared to other parenchymal axons within the same tissue; images displayed with Glow LUT **(B)**. Co-labeling of PGP9.5-EGFP^+/-^ (green) reporter mice demonstrated that NAN axonal varicosities contain the synaptic vesicle organizing protein, SYN1 (red) **(C)**, as well as synaptic vesicles (SV2, red) **(D)**. Varicose axons were observed surrounding unlabeled putative SCs (gaps between adjacent fibers; white arrows) in the parenchyma **(E)** and preceding many NANs **(C)**. SC identity was confirmed by SOX10 labeling; overlap of SOX10 (red) with DAPI (blue) within a nerve (green) leading to NAN **(F)**. White boxes were digitally magnified to aid in visualization. All images were captured on a Stellaris 5 confocal microscope. Lightning deconvolution was utilized to resolve axonal varicosities **(A-C**,**E)**. See Supplemental Figure S4 for single-color channels of each image.

Varicose axons were found throughout scWAT, with NANs characteristically displaying the greatest number of varicose axons (Figure 4B). To investigate whether synaptic transmission / neurotransmitter release could be occurring at theses nexuses, we immunostained whole PGP9.5-EGFP^+/-^ inguinal scWAT depots for the pre-synaptic vesicle organization protein, Synapsin I (SYN1) [28] (Figure 4C), and for the membrane glycoprotein SV2, which is localized to secretory vesicles [29] (Figure 4D). NANs were labeled by both characteristic pre-synaptic markers, which provided further evidence that these terminal structures are points of communication between the peripheral nervous system and single adipocytes. This is likely a site of release of the neurotransmitter norepinephrine, as NANs are largely labeled by TH (Figure 4A-B). However, the actual release and uptake of neurotransmitters at NANs has not yet been confirmed.

Varicose sympathetic axons were also visualized surrounding putative nerve-embedded SCs, as indicated by unstained gaps between adjacent fibers (Figure 4F, white arrows), similar to those previously shown to contain nuclei (Figure 2C) and confirmed to be SCs (Figure 3A). These same nerve-embedded SCs were frequently observed preceding NANs (Figure 4, white arrows and Supplemental Figure S4, white arrows). Co-staining of PGP9.5-EGFP^+/-^ inguinal scWAT revealed that these gaps between adjacent fibers were indeed SOX10+ SCs. To better characterize the NAN SCs, we stained whole inguinal scWAT from C57BL/6J mice for neural cell adhesion molecule (NCAM), a marker specific for nmSCs in adult mice [30] (Supplemental Figure S4D). However, we found that NCAM was not specific to nmSCs in scWAT and labeled all parenchymal nerves indiscriminately (Supplemental Figure S4E), including myelinated fibers (Supplemental Figure S4F).

Several similarities can be observed between the NAN and another peripheral nerve-termini, such as the well-characterized neuromuscular junction (NMJ) (Figure 3F). For both termini, the axon branches from peripheral bundles toward specific cellular targets, to form morphologically distinct connections. The axons leading to NMJs are myelinated [31], whereas the more distal pre-synaptic terminal is unmyelinated [32]. mSCs are found in/on the myelinated region of the NMJ axon, and nmSCs (important for maintaining synapses [32]) are found at the terminal, which are called terminal SCs (tSCs). In adipose, the NAN similarly branches from myelinated nerves with the terminal itself being unmyelinated, and like the NMJ, the NAN is immediately preceded by SCs.

### Fluorescence-activated cell sorting of SVF indicates the presence of two distinct SC populations

To determine both the presence and relative quantity of nmSC and mSC sub-populations, O4 and p75NTR were used as canonical markers. Previous literature indicated the use of O4 and p75NTR expression to positively select for SCs in rat sciatic nerve [33]. CD45 was used to negatively select for immune cells, after FSC/SSC gating on live singlet cells. O4 was used to positively select for mSCs, while O4 and p75NTR together were used to positively select for nmSCs [34]. The population of mSCs in scWAT was significantly higher compared to the nmSC population (Figure 5A) This was consistent with immunofluorescence labeling of SOX10+ SCs being associated primarily with myelinated structures in scWAT (Figure 3C). Since exercise has been shown to increase scWAT innervation [11], we aimed to investigate whether this intervention affects the relative distribution of SC populations in scWAT. Adult male BL6 mice were exercised for 7 days (caged with unrestricted running wheel access) or maintained sedentary (caged with locked running wheel). We observed no difference between exercised and sedentary SC populations in the scWAT of the two groups (Figure 5B). However, for both groups, the amount of mSCs trended higher compared to nmSCs (Figure 5B). Conversely, we have also previously shown that aging decreased innervation in scWAT [11]. Therefore, we examined whether aging alters SCs populations in scWAT. Following FACS of scWAT SVF from male BL6 mice across different age groups (4 months, 8 months, and 15 months), we again observed a higher amount of mSCs compared to nmSCs at 4 months (p=0.0011), 8 months (p=0.0017) and 15 months. At the latest age of 15mo, there was a loss of statistical significance (Figure 5C), possibly indicative of a relative decrease in myelinating SCs. Limitations in the FACS approach may preclude detailed phenotyping of SC subtypes in the tissue, including cell size exclusion and ‘stickiness’ of myelin to the adipocyte fraction.

**Figure 5:**
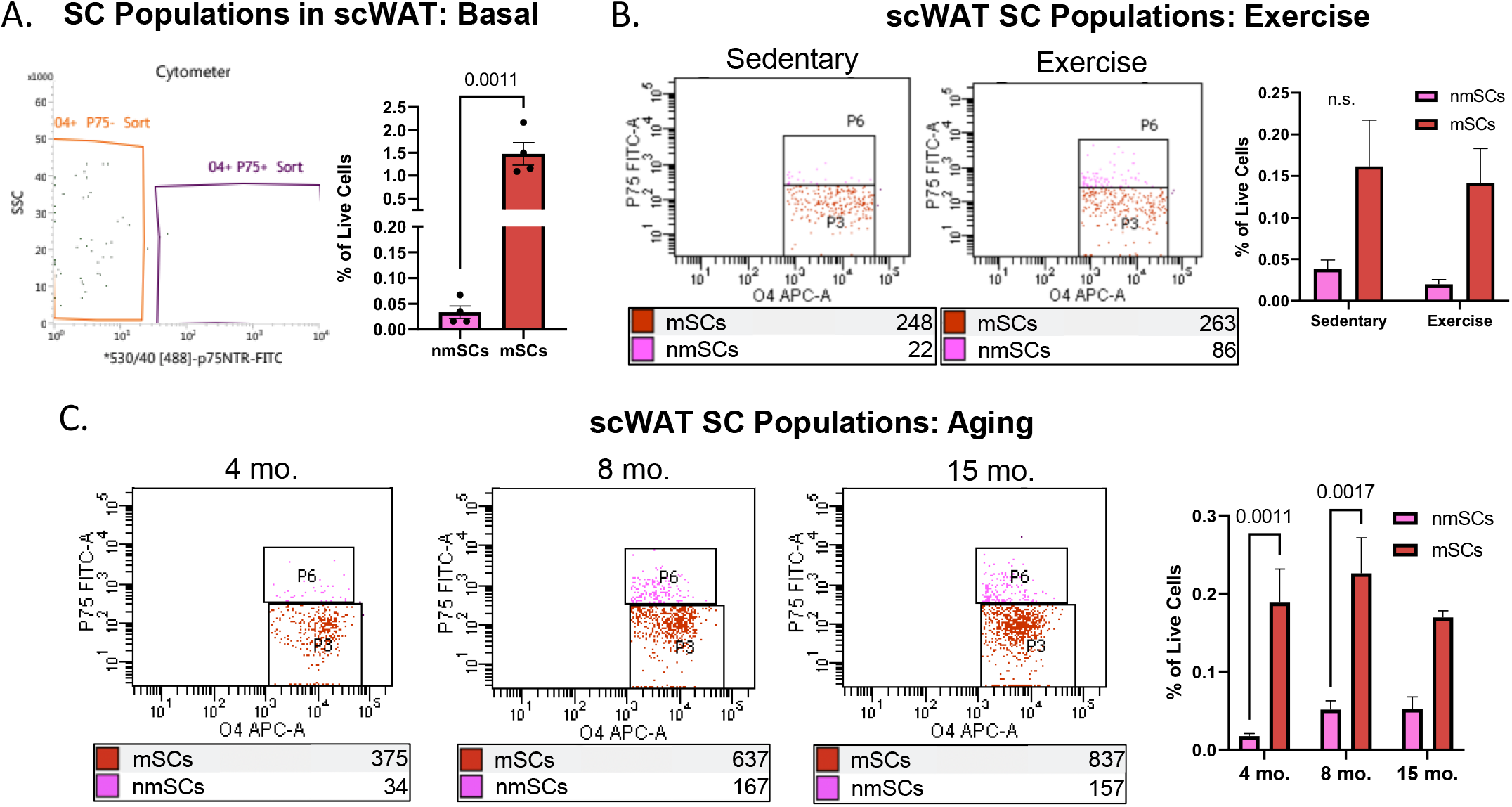
Fluorescence-activated cell sorting (FACS) and quantification of SCs in scWAT. Stromal vascular fraction (SVF) was isolated from inguinal scWAT depots excised from C57BL/6J mice and sorted into two distinct SC populations: CD45-O4+p75-, myelinating SCs (mSCs); and CD45-O4+p75+, non-myelinating SCs (nmSCs). mSCs and nmSCs were quantified as percent of live cells. Basal: 21-week-old male mice (N=4) **(A)**. Exercise: Male mice aged between 29 and 68 weeks were either given continuous access to run (Exercise, N=5) or had the running wheel locked in place (Sedentary, N=5) **(B)**. Aging: Male mice at three ages (4-months, N=5; 8-months, N=5; 15-months, N=5) **(C)**. Statistics: unpaired Student’s t-test **(A**) and one-way ANOVA with multiple comparisons **(B-C)**. Error bars are SEMs. P-values are as shown or are not provided when not significant (n.s.).

### Changes to SC phenotype with altered energy balance status

Adipose tissue innervation is highly plastic and imbalances in energy intake and energy expenditure have correlative impacts on tissue total innervation patterns, as well as sympathetic nerve activity [11; 35; 36], but the impacts of changing energy balance on adipose resident SCs have not yet been examined. We investigated gene expression changes in scWAT with obesity, aging, cold exposure, and exercise using six SC specific markers: the pan-SC markers *Sox10* [25; 26], oligodendrocyte marker 4 (*O4*) [37], and neurotrophin receptor p75 (*p75*) [37; 38]; the myelin-specific markers *Mpz* and *Krox20* [39; 40]; and the repair SC transcription factor *c-Jun* [17]. Because the associated changes in innervation status are likely to impact the total number of SCs, we also normalized gene expression to the pan-SC gene to investigate changes not confounded by changes in total SC number (Figure 6).

**Figure 6:**
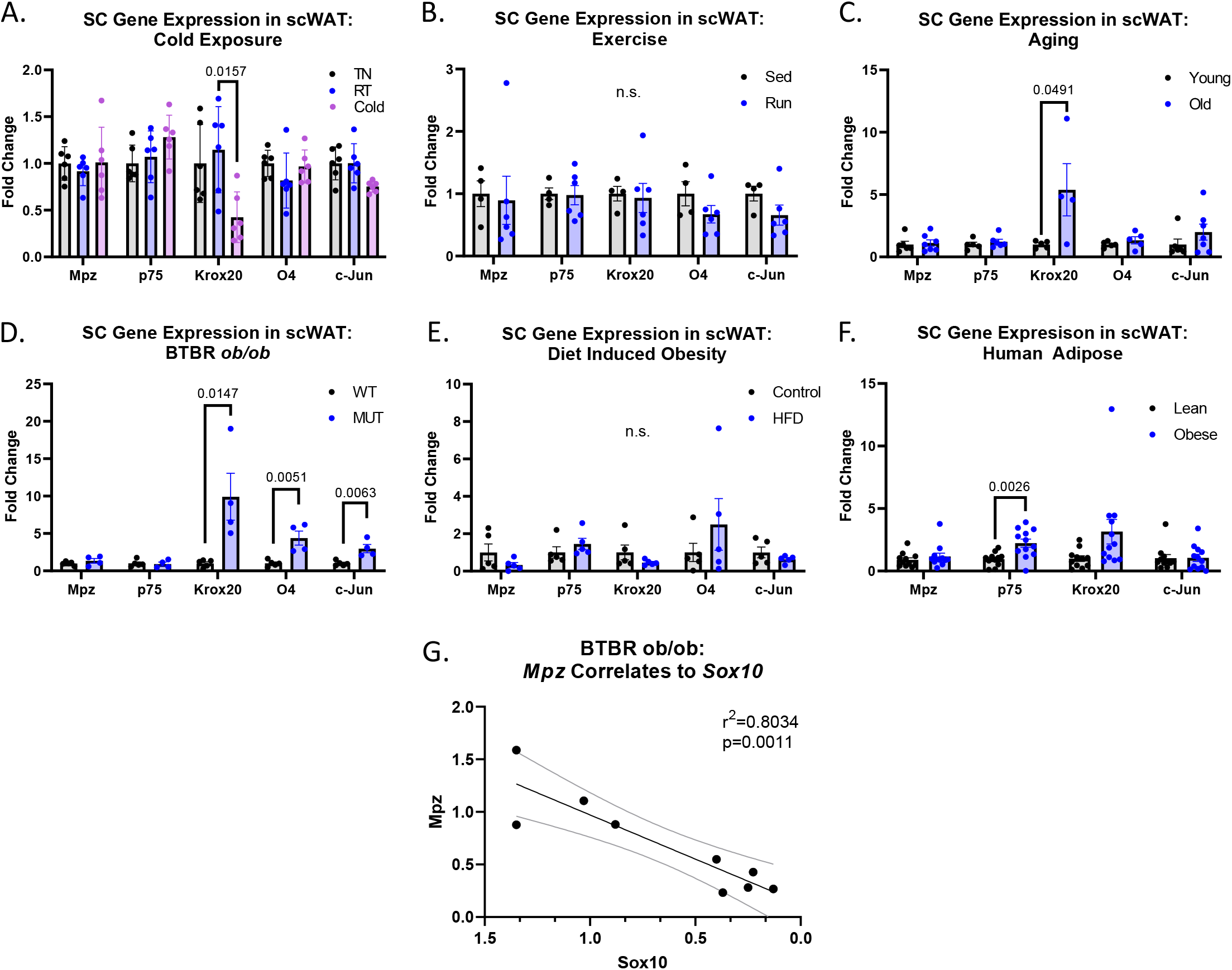
SC gene expression in scWAT with changing metabolic status. Relative gene expression measured by qPCR, normalized to total SCs (*Sox10*) and represented as fold change in ΔΔCt value **(A-F)**. Gene expression of scWAT from BL6 mice that were either housed at 30°C, thermoneutrality (TN, N=6); at 25°C, room temperature (RT, N=6); or at 5°C (Cold, N=6) **(A)**. Fold change is normalized to RT group. Gene expression of scWAT from BL6 mice that were exercised for 7-days or remained sedentary **(B)**. Mice were either given continuous access to running wheels (Exercise, N=6) or had the running wheel locked in place (Sedentary, N=4) **(B)**. Gene expression of scWAT from BL6 mice aged 15wks (Young, N=5) and 75wks (Old, N=4) **(C)**. Gene expression of scWAT excised from BTBR *ob/ob* (MUT, N=4) and wild-type littermates (WT, N=5) **(D)**. Gene expression of scWAT excised from BL6 mice fed either a 58% high fat diet (HFD, N=5) or chow (Control, N=5) **(E)**. Gene expression of scWAT biopsies excised from lean (BMI >30) and obese (BMI <25) human donors **(F)**. Linear regression of *Mpz* and *Sox10* foldchange normalized to housekeeper gene *Ppia* **(G)**. Statistics: one-way ANOVA with multiple comparisons **(A)** and unpaired Student’s t-test **(B-F)** and linear regression goodness of fit measured by R-squared and the significance of slope determined by F-test **(G)**. Error bars are SEMs. P-values are as shown or are not provided when not significant (n.s.). See Supplemental Figure S5 for all genes normalized to the housekeeper gene *Ppia*.

Cold exposure is a common means to increase sympathetic drive within adipose tissue and results in the tissue taking on a more metabolically favorable, energy expending phenotype with browning, increased sympathetic innervation, and non-shivering thermogenesis [35]. Male BL6 mice (N=6) were cold exposed at 5°C for 3 days and scWAT gene expression of SC markers was compared with mice housed at thermoneutrality (30°C) (N=6) or at room temperature (25°C) (N=6). Cold-exposed mice exhibited a downregulation in *Krox20* compared to mice housed at RT (p=0.0157) (Figure 6A).

Exercise promotes adipose tissue lipolysis through increased sympathetic drive and its demonstrated role in neuroplasticity [41], similar to cold exposure. Accordingly, we compared mice with access to voluntary exercise (running wheel cages; N=6) for 7 days with locked wheel caged sedentary littermates (N=4). Mice with access to running wheels displayed no significant differences in SC markers (Figure 6B) compared to sedentary controls (N=4).

Aging correlates with an increased prevalence of age-related neuropathy in the skin [42], the underlying muscle [32], and in scWAT [11]. We investigated the effects of aging on SC gene expression in scWAT by comparing male BL6 mice at 15 weeks old (N=5) to mice at 75 weeks old (N=4) (Figure 6C). Aged mice exhibited an upregulation in *Krox20* compared to the young controls (p=0.0491) (Figure 6C).

BTBR *ob/ob* (MUT) mice are leptin deficient (exhibiting an obese, diabetic, and neuropathic phenotype), with reduced innervation of scWAT [11; 43]. We compared relative gene expression of SC markers in MUT mice (N=4) to wild type BTBR (WT) littermate controls (N=5) (Figure 6D). MUT mice had a significant upregulation of *Krox20* (p=0.0147), *O4* (p=0.0051), and *c-Jun* (p=0.0063) (Figure 6D).

To investigate whether a diet-induced obesity (DIO) model would result in similar changes seen in obese BTBR *ob/ob* MUT mice, we measured relative gene expression in BL6 mice fed a high fat diet (HFD) for 19 weeks (N=5) compared to chow-fed controls (N=5) (Figure 6E). Interestingly, no significant changes were observed in SC marker gene expression, perhaps due to being less obese and neuropathic than the BTBR *ob/ob*. However, scWAT biopsies from male and female lean (N=11) and obese (N=12) human donors did display an upregulation in p75NTR (p=0.002) and a trending increase in Krox20 with obesity, which was more similar to what was observed in the BTBR *ob/ob* mice (Figure 6F).

Additionally, we normalized gene expression to the housekeeper gene *Ppia* to investigate what changes were occurring relative to the whole tissue (Supplemental Figure S5). We found that cold exposure resulted in down regulation of *Krox20* when compared to TN (p=0.0184) (Supplemental Figure S5A). There were no changes in gene expression with exercise or aging (Supplemental Figure S5B-C). Prominently, BTBR *ob/ob* MUT showed down regulation of *Sox10* (p=0.0076), *Mpz* (p=0.0089), and *p75* (p=0.0004), as well as upregulation of *Krox20* (p=0.0123) (Supplemental Figure S5D). A linear regression was performed looking at the relative fold change between *Sox10* and *Mpz* gene expression in BTBR *ob/ob* scWAT which found a strong correlation between the down regulation of *Mpz* with *Sox10* in scWAT (r^2^=0.8034, p=0.0011) (Figure 6G). Diet induced obesity in mice displayed no changes (Supplemental Figure S5E), but obese human scWAT showed an increase in *Krox20* gene expression (p=0.0456) (Supplemental Figure S5F).

When taking all data sets into consideration it becomes quite evident that pro-myelinating *Krox20* gene expression in scWAT is impacted by different metabolic states. Krox20 is consistently downregulated in scWAT under an energy expending metabolic state (cold exposure) and significantly upregulated in unfavorable metabolic states (aging and obesity). Importantly, these changes are consistent regardless of the reference gene.

#### BTBR *ob/ob* mice are neuropathic and show demyelination of small nerve fibers

BTBR *ob/ob* mice displayed the most changes in SC gene expression (Figure 6D) (Supplemental Figure S5D), and as such, we wanted to see if these changes were reflected in inguinal scWAT innervation and demyelination. Male and female BTBR *ob/ob* (MUT, N=4) and BTBR *+/+* wild-type littermates (WT, N=3) were aged to at least 12 weeks old (when they show an obese and neuropathic phenotype [11]). At the time of tissue collection, MUT mice had significantly greater body weight (p=0.0280), inguinal scWAT weight (p=0.0013), and subcutaneous adiposity (p=<0.0001) (Supplemental Figure S6A-C). Protein expression of whole inguinal scWAT lysates showed a decrease in sympathetic nerve activity (TH, p=0.0020) in MUT (Figure 7A) when compared to WT, as previously shown to correlate with a decrease in total innervation [11]. MUT mice also displayed a decrease in total SCs (SOX10) (p=0.0452) (Figure 7B) and a decrease in total myelination (MPZ) (p=0.0092) (Figure 7C).

**Figure 7:**
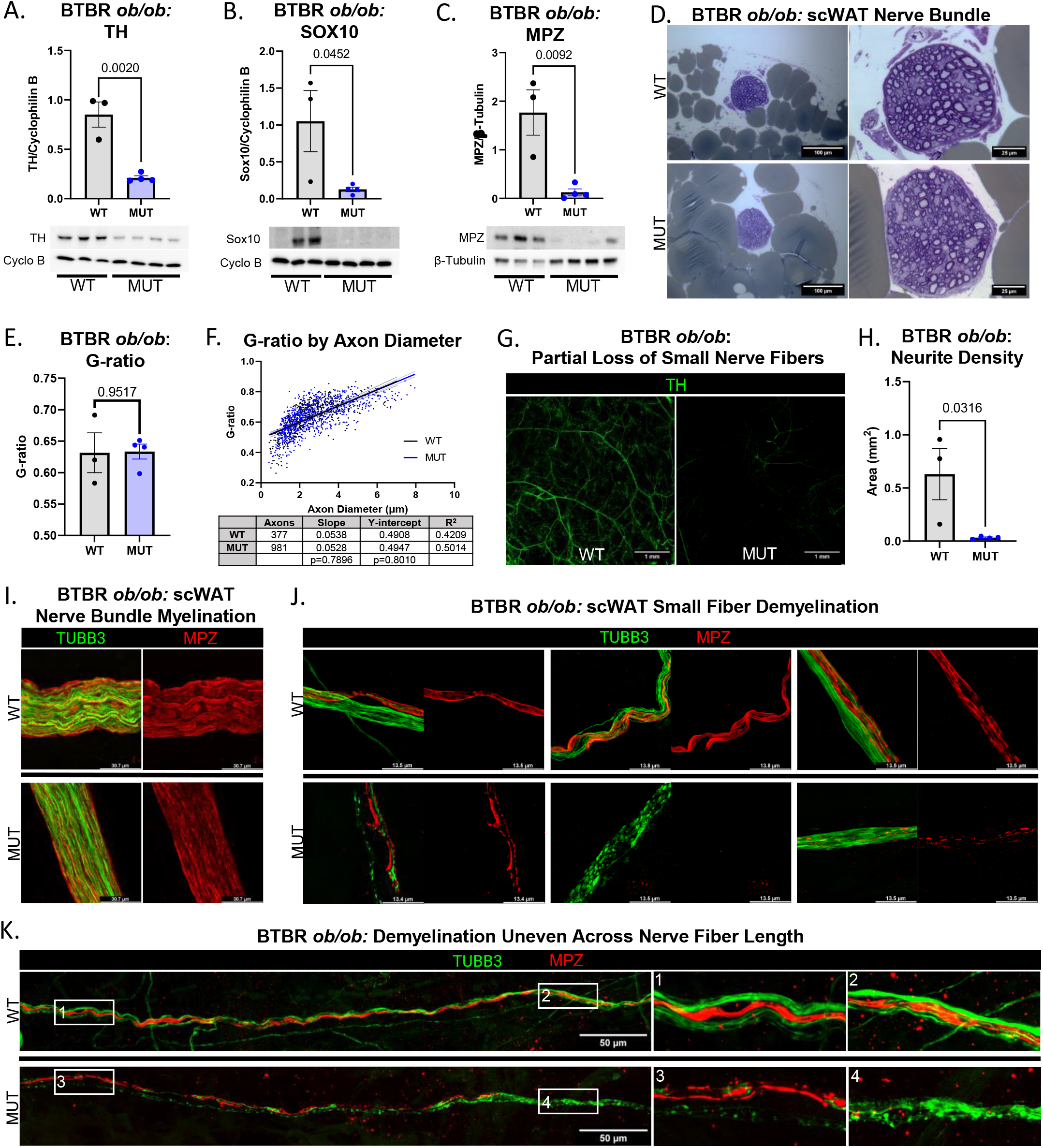
BTBR *ob/ob* mice present with a loss of small fiber sympathetic innervation in scWAT, accompanied by small fiber demyelination. Male and female BTBR *ob/ob* (MUT, N=4) and wild-type littermates (WT, N=3). Western blot protein analysis of TH **(A)**, SOX10 **(B)**, and MPZ **(C)** in inguinal scWAT. A consistent nerve bundle entering scWAT depot was excised from each mouse, cross-sectioned, and stained with toluidine blue **(D)**. G-ratio was measured for all myelinated axons in a nerve bundle and averaged for each mouse **(E)**. G-ratios were pooled for each group (WT, 377; MUT, 981) and plotted against axon diameter **(F)**. Linear regression values plotted in table below with slopes and Y-intercepts compared **(F)**. Whole mount immunostaining of sympathetic nerves in axillary scWAT were captured as Z-stacks (10 µm step size) with a 10X objective, tiled 5×5 (22.85 mm^2^ area), and maximum intensity projected **(G)**. Neurite density measured as fluorescence area in each 22.85 mm^2^ field of view **(H)**. Whole mount immunostaining of inguinal scWAT nerves (TUBB3, green) and myelin (MPZ, red) demonstrating myelinated nerve bundles **(I)** and demyelination of small nerve fibers in MUT mice **(J)**. Demyelination shown progressing along the length of comparable small fiber nerves overlying the subiliac lymph node (SiLN) in scWAT **(K)**. Numbers 1-4 correspond to adjacent images digitally magnified to aid visualization **(K)**. Brightfield images were captured on a Zeiss Axioskop microscope **(D)** and fluorescence images were captured on a Stellaris 5 confocal microscope **(G**,**I-K)**. Lightning deconvolution **(I-J)**. Statistics: unpaired Student’s t-test **(A-C**,**E**,**H)** and a linear regression **(F)**. Error bars are SEMs. P-values are as shown.

To gauge the scale at which demyelination was occurring, we started by consistently excising the same nerve bundle (Supplemental Figure S6D) entering the inguinal scWAT depot and analyzed nerve fiber myelination status by G-ratio. We observed no differences in the average G-ratio between WT and MUT mice (Figure 7E). There was also no change in G-ratio compared to axon diameter between both groups as determined by nearly identical linear regressions (Y-intercepts and slopes not significantly different) between WT and MUT mice (Figure 7F). With apparently no change in large bundle myelination, we turned our focus to the small fibers.

Whole axillary scWAT depots were excised (WT N=3; MUT N=4) and immunostained for TH which showed a loss of small TH+ fibers in the tissue parenchyma (Figure 7G), confirmed by quantifying neurite density (p=0.036) (Figure 7H). A similar decrease in scWAT innervation was observed in C57BL/6J *ob/ob* mice [44]. Additionally, we observed this same loss of small fibers in MUT inguinal scWAT, though with an insufficient sample size to draw conclusions (WT N=1; MUT N=1) (Supplemental Figure S6E). To investigate if the observed reduction in *Mpz* (Figure 6D) and MPZ (Figure 7C) was due to the overall reduced tissue innervation (Figure 7G-H) or a demyelination of the existing small nerve fibers we co-stained inguinal scWAT from BTBR *ob/ob* mice (WT N=3; MUT N=4) for the pan-neuronal marker, beta III tubulin (TUBB3) and MPZ. Immunostaining of tissue resident nerve bundles was consistent with our previous data, with no noticeable differences in bundle myelination (Figure 7I). However, many of the small fibers displayed a deterioration of the myelin sheath with the associated nerves taking on irregular shapes and a punctate appearance (Figure 7J). Because we could not differentiate between non-myelinated nerves, and those that had become completely demyelinated (though nerve irregularity was an indicator), we felt that we could not make definitive claims as to the ratio at which demyelination was occurring throughout each tissue. Nerves also did not show uniform demyelination along their length (Figure 7K); adding an additional layer of complexity in the neuropathy phenotype.

## DISCUSSION

Despite extensive research on SC development and injury responses within large nerve bundles such as the sciatic nerve, less is known about tissue-resident SCs and how unique environmental and metabolic cues may influence SC function. This is a significant gap in knowledge, given that chronic inflammatory demyelinating polyneuropathy (CIDP) is associated with metabolic conditions such as diabetes [45; 46] and the fact that demyelinating neuropathies may also be important in aging-related or idiopathic cases. Prior work has demonstrated the presence of SCs in brown adipose tissue [3; 5; 47], and scRNAseq studies have also reported the presence of SCs in white adipose [4; 6], with a recent study using SCs harvested from scWAT for regenerative therapies [8]. Regardless, it stands out that no studies have directly assessed the contributions of myelinated nerves and SCs to adipose tissue physiology. We sought to begin to fill this gap in understanding by investigating the tissue-resident SCs present in scWAT, how they contribute to myelinated axons in the tissue, and how myelination or SC phenotype may shift with metabolic state.

Our data provide conclusive evidence for myelinated and unmyelinated nerve subtypes within adipose, carried into scWAT by shared/mixed peripheral nerve bundles. Within these myelinated nerve bundles, we observed co-localized expression of both MPZ and MBP, demonstrating a role for both myelin proteins in adipose nerves. Of note, MBP+ myelination was observed in both CGRP+ sensory and TH+ sympathetic nerve fibers (Figure 2). This is contrary to statements in the available literature which state that post-ganglionic sympathetic nerves are largely unmyelinated [48]. Limitations in imaging or other experimental approaches may have led to this overly simplified conclusion. By contrast, previous work in rats had identified thinly myelinated sympathetic axons in the superior cervical ganglia and paravertebral chain ganglia [49], and more recently it was reported that MBP+ myelination may be protective against sympathetic denervation of the left ventricle in rhesus macaques [50]. Whereas these studies identified select populations of myelinated sympathetic nerves, we observed near complete overlap between TH and MBP within scWAT and found that the majority of sympathetic nerve fibers are at least thinly myelinated in the mouse scWAT tissue environment.

By contrast, we find that sensory (CGRP+) nerve fibers show greater heterogeneity in myelination. MBP and CGRP mark fibers (both thick and thin) entwined in nerve bundles together, further exemplifying the heterogenous nature of nerve bundles in adipose tissue. Sensory nerve fibers in the skin are also highly diverse; both functionally and characteristically. They can be either myelinated or non-myelinated, and when present, the myelin thickness corresponds to axon diameter and depends on several factors including the detected stimuli, whether the skin is glabrous or hairy, and even the dermal layer the nerve resides within [51]. Given our initial observations here, similar diversity in scWAT sensory fibers also exists and warrants further exploration.

The exact role of sympathetic myelination within adipose tissue is beyond the scope of this study but may strengthen the bidirectional communication between the CNS and adipose depots by allowing faster neural conduction, given myelin’s function as insulation for ionic movement in axons. Axonal diameter and myelin sheath thickness are directly related to the speed of signal conductance and resulting physiological functions [52]. Adipose nerves in the parenchyma also displayed heterogeneity in axon thickness and myelination, and may signify diversity in SNS functions in the tissue.

We were limited to assessing sympathetic and sensory myelination exclusively with anti-MBP labeling due to antibody host species cross-reactivity. This poses a significant caveat of these assessments, as we did observe less MPZ+ labeling around small neurovascular and parenchymal fibers (Figure 1), which tend to be sympathetic. As MPZ is the primary myelinating protein of the PNS, future studies utilizing an MPZ reporter mouse line would be important for clarifying this matter and are underway in our laboratory currently.

The presence of nerve termini in scWAT, unique from the majority of innervation observed across the tissue architecture, is intriguing, but ultimately provides more questions than it does answers. It is still unclear if a true synapse is being formed or what the function is of these terminal endings that only form connections with a subset of mature adipocytes. Many of the parenchymal nerves in scWAT are varicose, and it was shown in BAT that sympathetic axonal varicosities can make direct contact with adipocytes [3]. We have now shown that NANs house synaptic vesicles in axonal varicosities and likely function as a pre-synapse terminal releasing sympathetic neurotransmitters on effector adipocytes. However, a true post-synaptic junction on adipocytes has yet to be observed or described, despite the presence of post-synaptic proteins expressed in adipose tissue, such as PSD95 [11]. This may instead be similar to other autonomic neuroeffector junctions which are characterized by varicose nerve fibers that release neurotransmitters onto effector cells that do not contain post-synaptic specialization but do have neurotransmitter receptors on their cell membrane [27]. As mentioned, many of the sympathetic nerves in scWAT parenchyma were varicose suggesting that they may be largely releasing neurotransmitter and neuropeptide along the length of their axons (‘en passant’) onto contacted adipocytes. NANs would accordingly serve a specialized function requiring more targeted synaptic release versus the diffuse release that occurs en passant. The distinguishing characteristics between an adipocyte which forms a nexus and one that does not is still a mystery, but is supported by the vast literature describing numerous subtypes of scWAT mature adipocytes. The mechanisms that induce nexus formation also remain to be determined.

By characterizing the glia associated with NANs, such as the SCs that appear similar to the terminal SCs observed at the NMJ, we hoped to further develop our understanding of NANs as well as the important functions SCs serve that are unique to adipose tissue. NCAM labeling indicated that these NAN terminals are unmyelinated, but this was undermined by the NCAM antibody’s apparent lack of binding specificity within a tissue environment. The presence of SCs immediately preceding the NAN suggests the alternative; that the axons leading to each NAN are myelinated, as we have demonstrated that scWAT SCs are mostly myelinating and tend to be localized to myelinated structures. Regardless, the consistency with which SCs are localized to NANs hints to their importance for maintaining and/or eliciting these connections. By drawing comparisons with the NMJ (a well described peripheral nerve terminal) we hoped to tease out potential functional similarities shared between the supporting glia at each terminal. tSCs are crucial for maintaining NMJs so we hypothesized that tSCs would be present at the terminal junction of the NAN as well. However, the terminal SCs of the NAN were different than the NMJ, in that they were only observed in the preceding axon leading to the NAN and not at the terminal itself. These SCs still may be functioning as tSCs in synapse maintenance and nerve repair, but hints that the long-term maintenance provided by abundant tSCs as required by the NMJ, may not be required by NANs. This could suggest that NANs are highly plastic and do not form life-long connections.

The frequency of nerve myelination within adipose tissue, and presence of both mSCs and nmSCs (Figure 5) subtypes, demonstrates a role for SCs in adipose nerve maintenance and function and makes them a potential target of dysregulation during adipose neuropathy. We have shown that the scWAT of BTBR *ob/ob* MUT mice became neuropathic, with observed demyelination of small fibers in the tissue. Additionally, these obese mice also underwent the most changes in SC-related genes across changing energy balance states, specifically exhibiting upregulation of *Krox20* (Figure 6). Krox20 has been identified as the main regulator of SC myelination during the pro-myelination phase, and plays a key role in interacting with Sox10 during the formation of nodes of Ranvier. Krox20 also acts as a transcription factor for the production of MPZ [37; 53; 54], thus emphasizing Krox20’s importance in myelination of axons [39; 55; 56].

We also observed a significant upregulation of Krox20 in BL6 mice with age and in humans with obesity (Figure 6). Interestingly, in each instance, *Krox20* increased without a correlative increase in *Mpz* expression (Figure 6). Moreover, BTBR *ob/ob* mice displayed reduced myelination (Figure 7) alongside reductions in *Mpz* when normalized to *Ppia*, but this difference disappeared when normalized to *Sox10* indicating that the downregulation of *Mpz* was directly correlated to total SC *Sox10* expression. This was confirmed by running a linear regression of both genes which correlated the downregulation of *Mpz* with a downregulation of *Sox10* (Figure 6). We hypothesize that upregulation of *Krox20* may be a compensatory response of adipose mSCs to increase myelin production in neuropathic environments. Additionally, an increase in Krox20 expression would be expected if mature SCs were transdifferentiating into repair SCs [16] in response to obesity or age-related neuropathy. The observed decrease in Krox20 expression may then indicate an impaired ability to produce repair SCs in neuropathic states. Further research is needed to identify environmental signals that may be preventing functional myelination in these models despite robust Krox20 activation, or whether Krox20 may be functioning in other ways in the tissue.

A hallmark feature of SCs is their ability to shift towards a repair phenotype in response to injury, releasing neurotrophic factors such as BDNF and glial cell line-derived neurotrophic factor (GDNF). Previous studies have demonstrated the decreased ability of aged mice to regenerate nerves post-injury when compared to their younger counterparts as a result of decreased c-Jun expression [17], and neuropathic diseases such as CIDP are characterized by a loss of SC plasticity [57]. Notably, qPCR of *c-Jun* in scWAT revealed almost no difference between young and old mice while displaying upregulation in obese BTBR *ob/ob* mice (Figure 6). In order to promote Wallerian degeneration, c-Jun is an inhibitor of myelination [58], and recent studies found that SC mitochondrial dysfunction may drive c-Jun expression that contributed to demyelination [59]. The upregulation of *c-Jun* in obese BTBR *ob/ob* mice may be contributing to the demyelination of small fibers, or alternatively, as a means for repairing the neuropathy that has occurred.

We have previously shown that obese BTBR *ob/ob* mice display neuropathy in scWAT [11; 43] and others have demonstrated a specific decline in scWAT neurovascular innervation [11; 43]. Here we have provided evidence that this neuropathy likely begins with the small parenchymal nerve fibers and is associated with deteriorating myelin sheaths. It is still unclear if demyelination is causing the axonopathy. We noted that MPZ did not label many small parenchymal nerve fibers in scWAT (which are mostly sympathetic). Interestingly, in BTBR *ob/ob* mice we observed deterioration of myelin sheaths labeled with MPZ as well as a decrease in small sympathetic fibers. This suggests that there is axonopathy of the sympathetic fibers that may be independent from the demyelination. Whether or not the neuropathy effects both sensory and sympathetic nerves equally is also unclear. The downregulation of *Mpz* correlated with a downregulation of *Sox10* and decreased SOX10 protein expression, but whether this is a loss of just mSCs, or both mSCs and nmSCs is unclear. Finally, one could imagine that therapies capable of preventing or reducing the loss of adipose tissue resident SCs may protect from nerve loss and demyelination, ultimately contributing to healthier metabolism and energy balance, especially in metabolic disease states like obesity and diabetes

In conclusion, we have now characterized SCs in white adipose tissue, their association with myelinated and non-myelinated nerves, and localization to synaptic vesicle-containing terminal nerve structures, or NANs. Most importantly we have provided evidence that obesity/diabetes-related adipose neuropathy is concurrent with small fiber demyelination, loss of small fiber innervation, and a decrease in SCs - together illustrating the importance of SCs in the maintenance of adipose tissue peripheral nerves and thereby healthy metabolism.

## ACKNOWLEDGEMENTS

The authors wish to thank Morganne Robinson and Joshua Havelin for technical assistance, as well as FACS services at the Jackson Laboratory (Will Schott), The Ohio State University Wexner Medical Center Flow Cytometry Core, and Nationwide Children’s Hospital (Dave Dunaway). We would also like to thank Robert Burgess at the Jackson Laboratory for technical advice.

## AUTHOR CONTRIBUTIONS

JWW, GG, EP, and MB wrote the manuscript, designed experiments, and analyzed data. JRT processed and imaged tissues for brightfield microscopy. MFP, SRS, and LMS conducted the human tissue collection and provided samples. KLT wrote the manuscript, designed experiments, analyzed data, conceived of and oversaw the project, and serves as lead contact for this manuscript.

## GUARANTOR STATEMENT

Kristy L. Townsend is takes responsibility for the research presented in this manuscript.

