## Supplemental Table 1 and Supplemental Figure Legend for "The Presence of Myelinated Nerves and Schwann Cells in White Adipose Tissue: Proximity to Synaptic Vesicle Containing Nerve Terminals and Potential Role in BTBR *ob*/*ob* Demyelinating Diabetic Neuropathy"

**SUPPORTING INFORMATION:**

**Supplemental Table 1. qPCR primer sequences**.

| Gene | Target | Forward Sequence 5' → 3' | Reverse Sequence 5' → 3' |
|---|---|---|---|
| *c-Jun* | Mouse | TGG CAT CAC CAC TAC AC | TCT GGC TAT GCA GTT CAG CC |
| *c-Jun* | Human | TAA CAG TGG GTG CAA ACT CA | TTT TTC TCT CCG TCG CAA CT |
| *Krox20* | Mouse | GCC CCT TTG ACC AGA TGA AC | GGA GAA TTT GCC CAT GTA AGT G |
| *Krox20* | Human | GAC CAT CTT TCC CAA TGC CG | TTT CTA GGT GCA GAG ACG GG |
| *Mpz* | Mouse | GTC AAG TCC CCC AGT AGA A | GTC AAG TCC CCC AGT AGA A |
| *Mpz* | Human | AAG TGC CAA CTA GGT ACG GG | CAT AGC ACT GAG CCT CCT CT |
| *O4* | Mouse | TGC CTG GGG AAA TCA GTC AT | AGT TCG TCC ATT TTT CGG CAG |
| *Ppia* | Mouse | GGC AAA TGC TGG ACC AAA C | CAT TCC TGG ACC CAA AAC G |
| *Ppia* | Human | GTC AAC CCC ACC GTG TTC TTC | TTT CTG CTG TCT TTG GGA CCT TG |
| *p75ntr* | Mouse | TGC CTG GAC AGT GTT ACG TT | ACA GGG AGC GGA CAT ACT CT |
| *p75ntr* | Human | CCT ACG GCT ACT ACC AGG ATG AG | TGG CCT CGT CGG AAT ACG |
| *Sox10* | Mouse | AGA TCC AGT TCC GTG TCA ATA A | GCG AGA AGA AGG CTA GGT G |
| *Sox10* | Human | TCA TCC CTT CAA TGC CCC CT | TGC GTC TCA AGG TCA TGG AGG |

**Supplemental Figure S1: Single-color channels related to figure 1.** Intact scWAT depots were excised from PGP9.5-EGFP$^{+/-}$ (green) reporter mice and immunolabeled for myelin with MPZ and MBP. Related to Figure 1A; MPZ (red) **(A)**. Related to Figure 1B; MBP (red) **(B)**. Related to Figure 1C; MBP (red), MPZ (blue) **(C)**. Images were captured on Stellaris 5 confocal microscope.

**Supplemental Figure S2: Single-color channels related to figure 2.** Intact scWAT depots were excised from PGP9.5-EGFP$^{+/-}$ (green) reporter mice and immunolabeled for myelin with MBP (blue) and either sympathetic nerves (TH) or sensory nerves (CGRP). Related to Figure 2A; TH (red) **(A)**. Related to Figure 2B; CGRP (red) **(B)**. Related to Figure 2C; Top row (TH, red), middle and bottom rows (CGRP, red) **(C)**. Images were captured on Stellaris 5 confocal microscope.

**Supplemental Figure S3: Additional co-labeling of SOX10 and single-color channels related to figure 3.** Intact scWAT depot excised from female PGP9.5-EGFP$^{+/-}$ (not shown) mouse labeled for SOX10 (cyan) and imaged as a tiled $Z_{max}$ projection of the whole tissue **(A)**. High magnification images of female PGP9.5-EGFP$^{+/-}$ (yellow) mouse labeled for SOX10 (cyan) and DAPI (red) which revealed that SOX10 had off-target labeling of mammary ducts and/or lymphatics **(B)**. Digital cross section (bottom row) **(B)**. Single-color channels related to figure 3C; DAPI (white), PGP9.5-EGFP$^{+/-}$ (green), SOX10 (red), and MBP (blue). Images were captured on Stellaris 5 confocal microscope.

**Supplemental Figure S4: Additional labeling of neuro-adipose nexuses (NANs), NCAM off-target labeling, and single-color channels related to figure 4.** NANs in BL6 mouse inguinal scWAT labeled for TH and imaged without adipocyte autofluorescence **(A).** Single color channels of PGP9.5-EGFP$^{+/-}$ (green) mice related to Figure 4C (SYN1, red) **(B)**, related to Figure 4D (SV2, red) **(C)**, and related to figure 4F (DAPI, blue; SOX10, red) **(D)**. Intact scWAT depots excised from BL6 mice were immunostained for NCAM (orange), a canonical non-myelinating Schwann cell (nmSC) marker, which labeled NANs **(E)**. PGP9.5-EGFP$^{+/-}$ (green) mice immunostained for NCAM (red) found that in addition to SCs, NCAM labeled parenchymal nerve fibers **(F)**, some of which were myelinated (MBP, blue) **(G)**. Diagram of a NAN and a neuromuscular junction (NMJ) demonstrating similarities and differences of both peripheral nerve terminals in respect to myelination and SCs. Fluorescence images are provided of both terminal structures for reference: NAN in scWAT labeled with PGP9.5-EGFP$^{+/-}$ reporter; NMJ labeled with 2H3 and SV2 for pre-synaptic terminal, α-bungarotoxin (BTX) for post-synaptic terminal, MPZ for myelination, and DAPI for nuclei **(G)**. Circles outline each nexus; white arrows point to presumed SCs indicated by gaps between adjacent fibers **(A,C-D,G)**. Images captured on Stellaris 5 confocal microscope **(A-D,F-H)** and Eclipse E400 epifluorescence microscope **(E).**

**Supplemental Figure S5. SC gene expression in scWAT with changing metabolic states; normalized to *Ppia*.** Relative gene expression measured by qPCR, normalized to the housekeeper gene *Ppia* and represented as fold change in ΔΔCt value. Gene expression of scWAT from BL6 mice that were either housed at 30$^{o}$C, thermoneutrality (TN, N=6); at 25$^{o}$C, room temperature (RT, N=6); or at 5$^{o}$C (Cold, N=6) **(A)**. Fold change is normalized to RT group. Gene expression of scWAT from BL6 mice that were exercised for 7-days or remained sedentary **(B)**. Mice were either given continuous access to running wheels (Exercise, N=6) or had the running wheel locked in place (Sedentary, N=4) **(B)**. Gene expression of scWAT from

BL6 mice aged 15wks (Young, N=5) and 75wks (Old, N=4) **(C)**. Gene expression of scWAT excised from BTBR ob/ob (MUT, N=4) and wild-type littermates (WT, N=5) **(D).** Gene expression of scWAT excised from BL6 mice fed either a 58% high fat diet (HFD, N=5) or chow (Control, N=5) **(E)**. Gene expression of scWAT biopsies excised from lean (BMI >30) and obese (BMI <25) human donors **(F)**. Statistics: one-way ANOVA with multiple comparisons **(A)** and unpaired Student's t-test **(B-F)**. Error bars are SEMs. P-values are as shown or are not provided when not significant (n.s.).

**Supplemental Figure S6: BTBR *ob/ob* weight and adiposity, immunostaining, and single-color channels related to figure 7.** Male and female BTBR *ob/ob* (MUT, N=4) and +/+ wild-type littermates (WT, N=3). Body weight **(A)** and scWAT weight **(B)** recorded at time of tissue collection. Subcutaneous adiposity was measured as (scWAT weight / body weight) **(C)**. Image captured through the eyepiece of a dissecting microscope to show the location of the nerve bundle that was consistently excised for g-ratio measurements **(D)**. An Intact scWAT depot from a WT and a MUT was immunostained for TH **(E)**. Black arrows point to nerve bundle entering scWAT, White arrow points to the subiliac lymph node (SiLN) **(E)**. Single-color channels of whole mount scWAT immunostained for TUBB3 (green) and MPZ (red) **(F)**. Fluorescence images captured on Stellaris 5 confocal microscope. Statistics: unpaired Student's t-test **(A-C)**. Error bars are SEMs. P-values are as shown.
