## Supplemental Figures for "The Presence of Myelinated Nerves and Schwann Cells in White Adipose Tissue: Proximity to Synaptic Vesicle Containing Nerve Terminals and Potential Role in BTBR *ob*/*ob* Demyelinating Diabetic Neuropathy"

Supplemental Figure S1: Single-color channels related to figure 1.

A.

**Single-color Channels of Figure 1A**

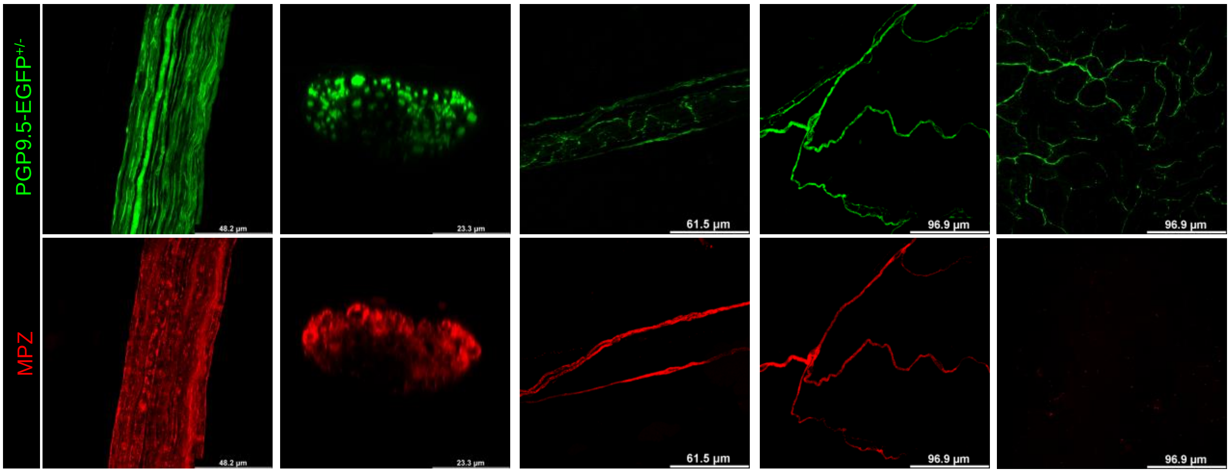


B.

**Single-color Channels of Figure 1B**

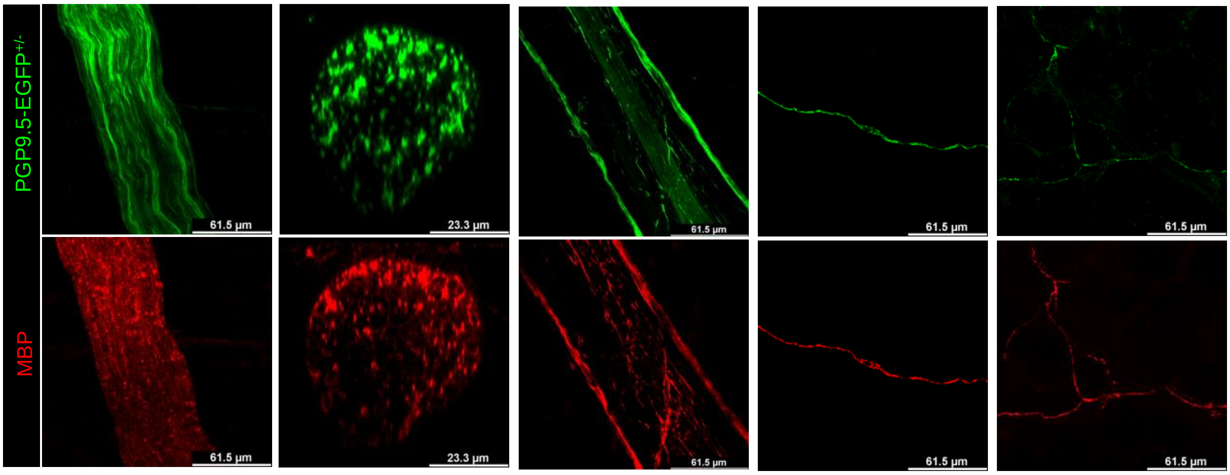


C.

**Single-color Channels of Figure 1C**

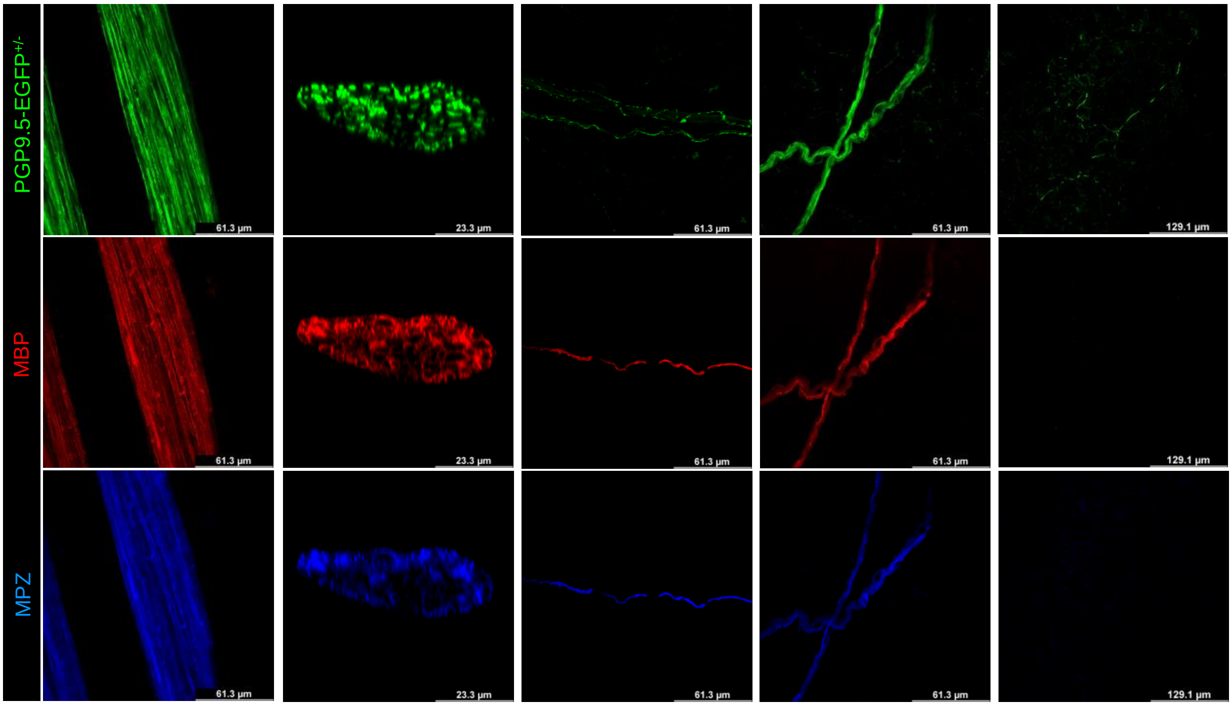

Supplemental Figure S2: Single-color channels related to figure 2.

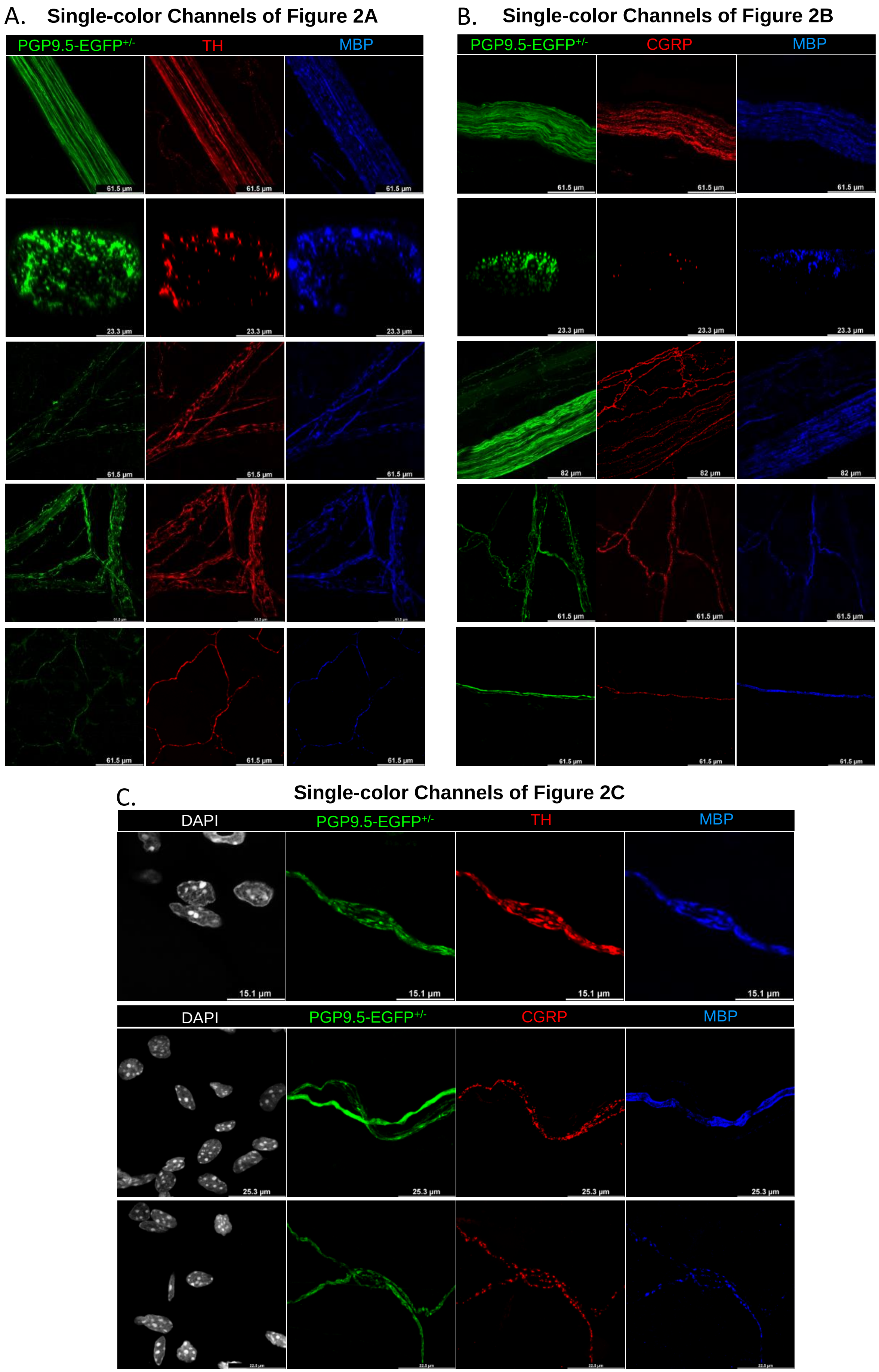

Supplemental Figure S3: Additional co-labeling and single-color channels related to figure 3.

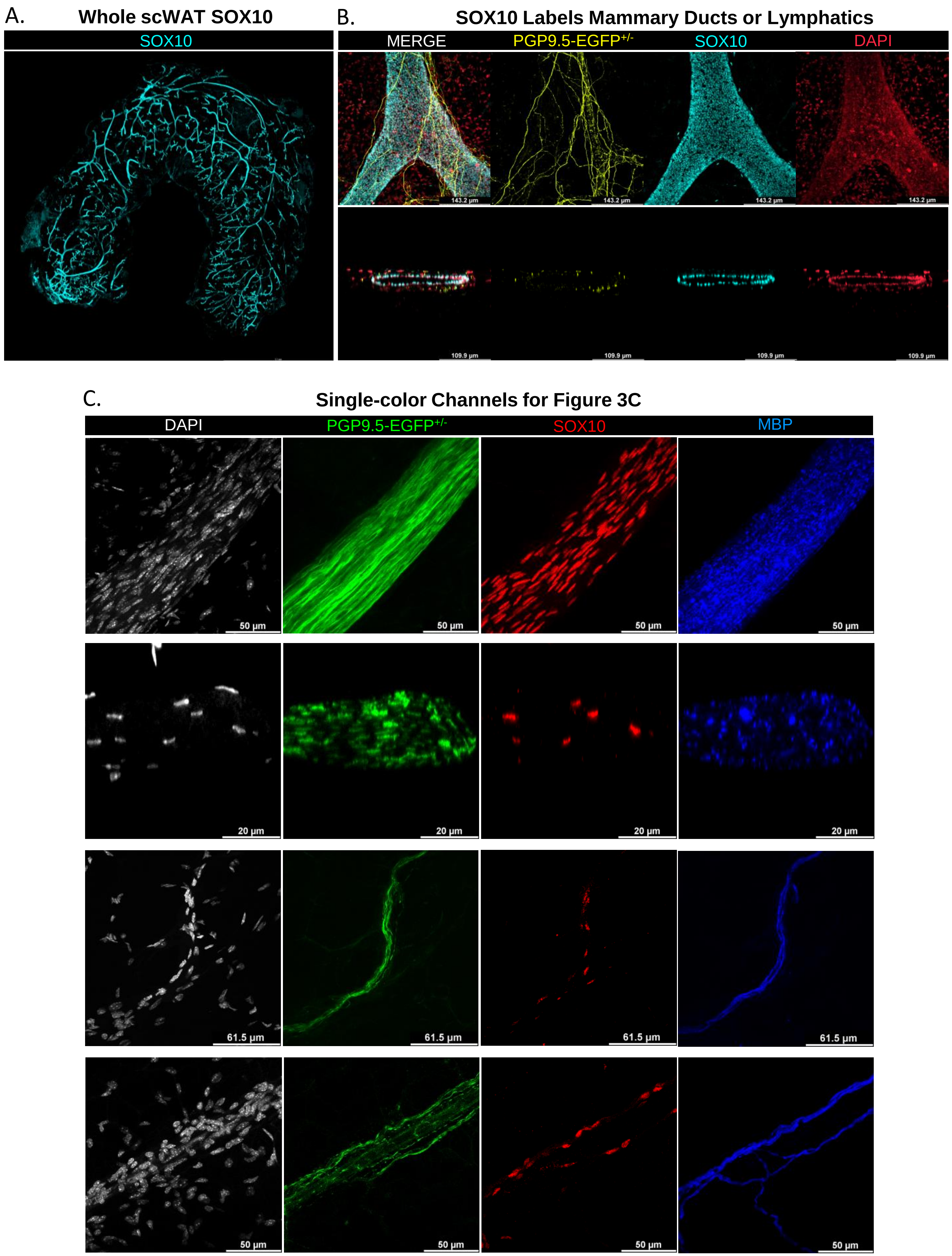

Supplemental Figure S4: Additional labeling of neuro-adipose nexuses (NANs), NCAM off-target labeling, and single-color channels related to figure 4.

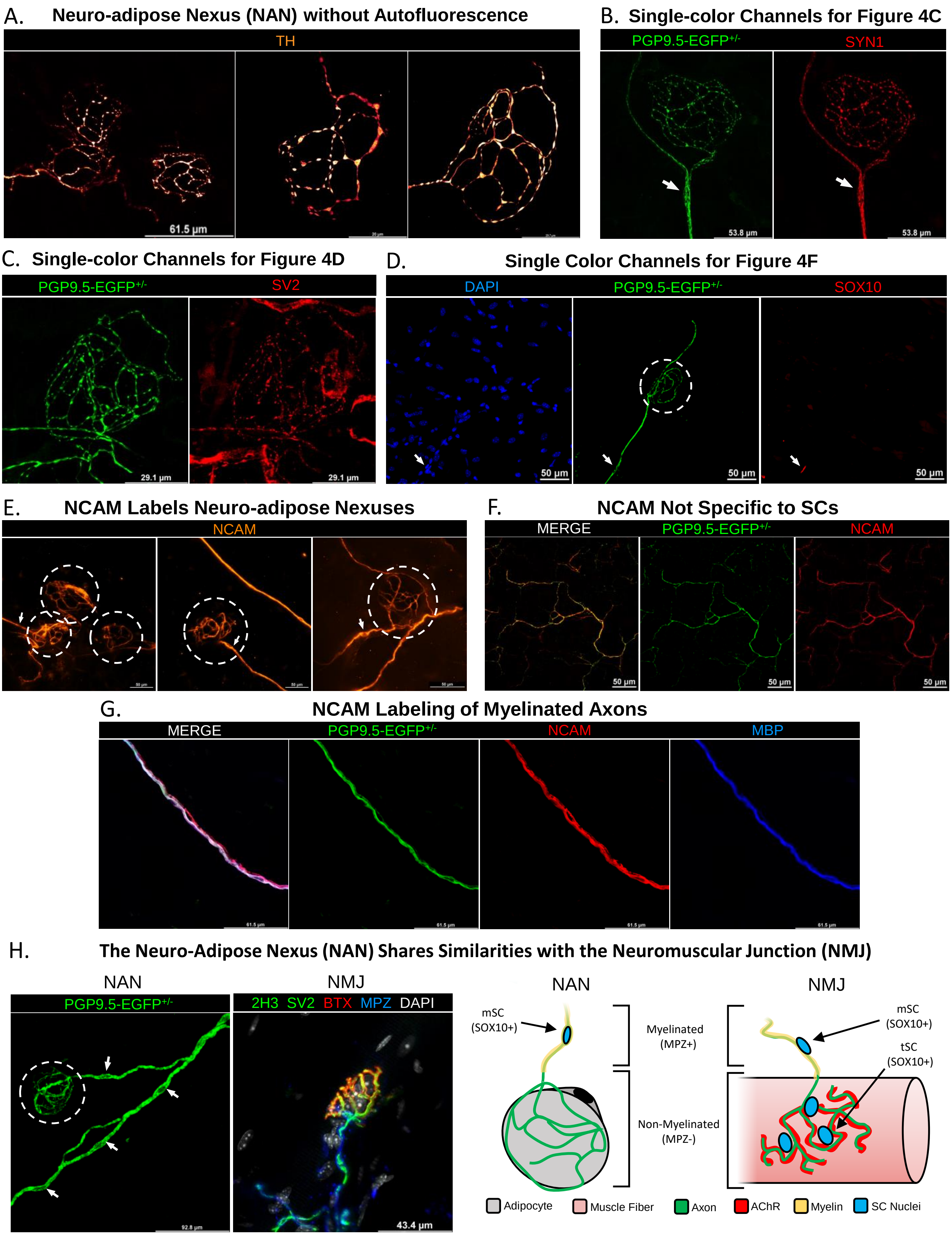

Supplemental Figure S5: SC gene expression in scWAT with changing metabolic status; normalized to *Ppia*.

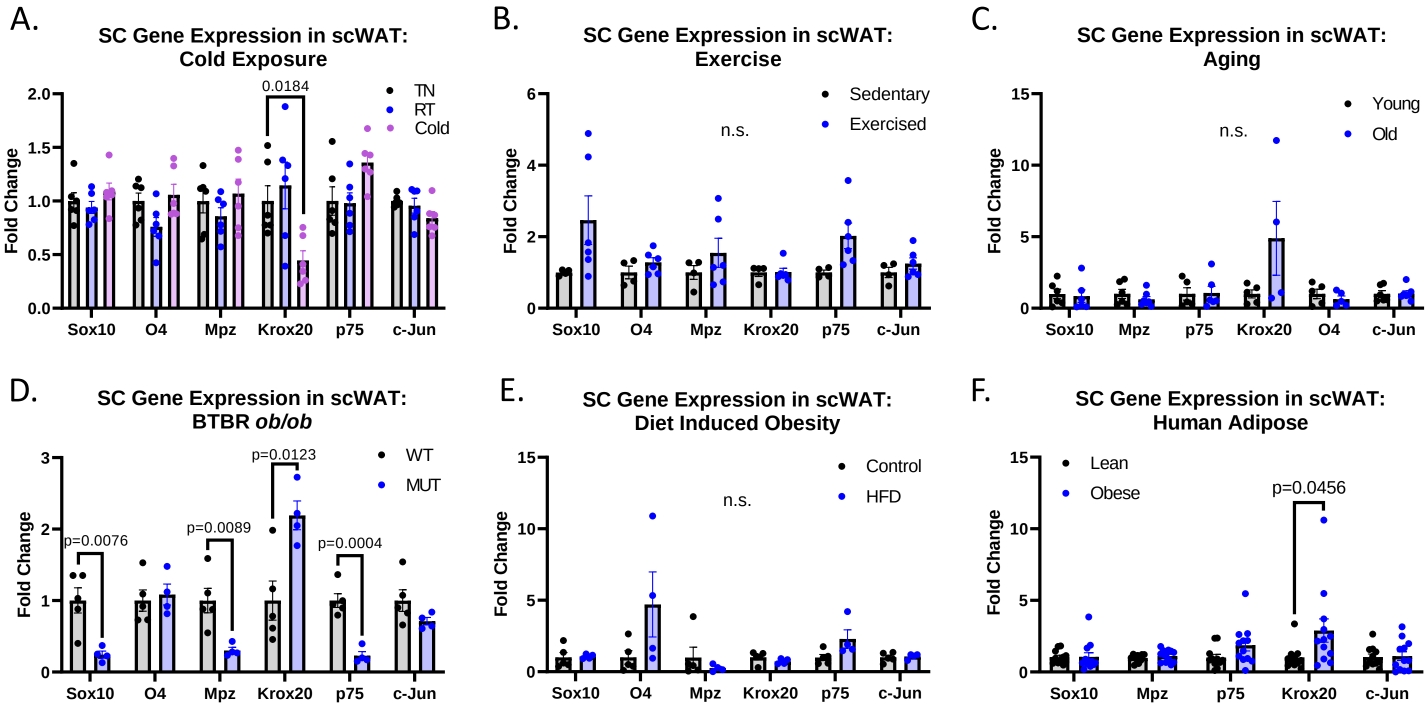

Supplemental Figure S6: BTBR *ob/ob* weight and adiposity, immunostaining, and single-color channels related to figure 7

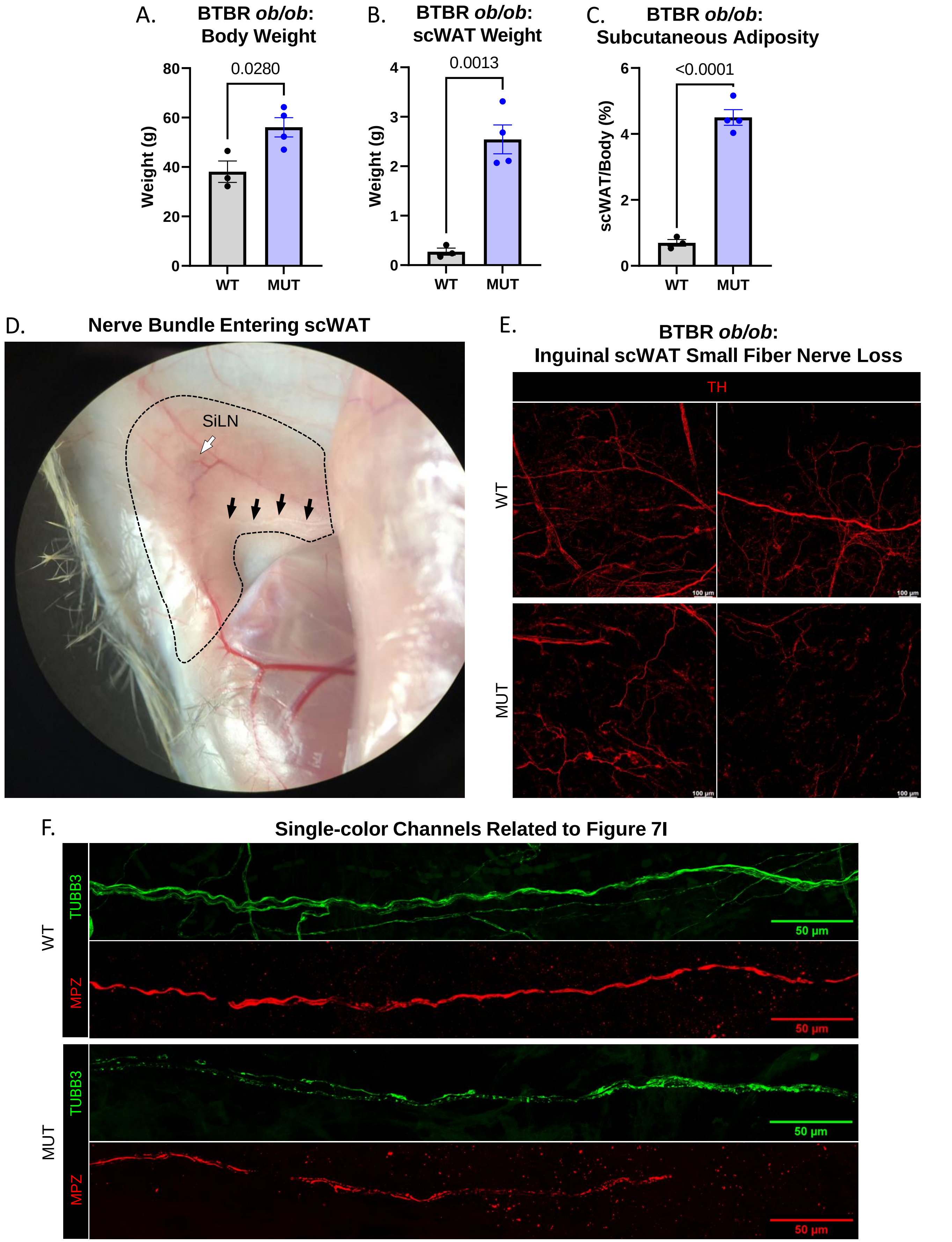
